# Dorsal fins are not universal stabilizers in cetaceans: limited yaw effects and flipper-coupled roll stability

**DOI:** 10.64898/2026.04.15.718588

**Authors:** Taro Okamura, Masateru Maeda, Futaba Nishimura, Ken Yoda

## Abstract

Cetacean dorsal fins are traditionally regarded as vertical stabilizers for yaw and roll, yet marked variation in fin area and position suggests that this function is not universal. We combined computational fluid dynamics simulations of five cetacean species with comparative analyses of dorsal fin and flipper morphology across 81 species to test whether variation in dorsal fin morphology reflects the evolution of hydrodynamic stabilization. Yaw-destabilizing moments were dominated by the trunk regardless of flipper posture, whereas dorsal fins were generally too small and too close to the center of rotation to provide substantial static yaw restoration. In contrast, dorsal fins influenced roll stability in concert with extended anhedral flippers. Although dorsal fins were likely present in the common ancestor of extant cetaceans, strong dorsal fin-mediated roll stabilization was largely restricted to oceanic bite-feeding delphinids, in which rapid evolutionary enlargement of the dorsal fin and persistently extended anhedral flippers likely enhanced roll stability. In most other lineages, roll stability could be maintained by flipper posture alone despite small dorsal fins. These results recast the cetacean dorsal fin not as a universal stabilizer, but as a lineage-specific roll stabilizing structure whose function emerges through mechanical coupling with the flippers.

## Introduction

The dorsal fin of cetaceans represents a prominent morphological novelty and a hallmark of convergent evolution among aquatic vertebrates. Despite its marked morphological diversity and long-standing use in species identification^1–4^, its functional significance remains unclear. In aquatic vertebrates, fins generally contribute to propulsion and stability^5–9^, and variation in fin morphology often reflects differences in swimming performance associated with lifestyle^10–14^. During secondary adaptation to aquatic environments, cetaceans evolved three principal appendages: flippers, a dorsal fin, and a caudal fin^15–18^. The jointless dorsal fin has long been interpreted as a fixed vertical stabilizing surface that contributes to yaw and roll stability, analogous to the vertical stabilizer of an aircraft or the keel of a ship^7,8,19–24^. This interpretation appears to be consistent with the relatively large dorsal fins observed in several small odontocetes and in killer whales (*Orcinus orca*). In contrast, baleen and beaked whales possess relatively small, posteriorly positioned dorsal fins^15,25^. Furthermore, some species including belugas (*Delphinapterus leucas*) and finless porpoises (*Neophocaena* sp.) lack a distinct dorsal fin^2,3,26^. If dorsal fin morphology was governed by strong and uniform functional constraints, convergence in fin area and position would be expected across cetaceans. Instead, this pronounced interspecific variation suggests that the dorsal fin does not function as a universal vertical stabilizer. This issue is particularly important in cetaceans because, unlike cartilaginous and ray-finned fishes and Mesozoic marine reptiles such as ichthyosaurs and mosasaurs, they lack a vertically oriented caudal fin.

The hydrodynamic roles of cetacean fins, including the dorsal fin, have been investigated using experimental and computational approaches^27–32^. However, most studies have examined fins in isolation, leaving several key questions unresolved: how much the dorsal fin contributes to whole-body static stability, whether this contribution is associated with flipper morphology and posture, and whether these relationships reflect ecological diversity. In particular, flippers can also contribute to yaw and roll stability, sugeesting a hydrodynamic functional overlap with the dorsal fin^6,33,34,35^. Although flippers may be folded against the trunk during straight-line swimming to reduce drag^25,36^, small cetaceans hold their flippers extended at pronounced anhedral angles, which may affect whole-body stability^36,37^. Interspecific variation in flipper anhedral angle has also been associated with the presence or absence of a dorsal fin^36^. These observations suggest that the hydrodynamic function of the dorsal fin must be evaluated within an integrated whole-body framework that incorporates flipper morphology and posture.

An integrated hydrodynamic perspective on dorsal fin and flipper function may provide a mechanistic basis for ecological diversification in cetaceans, because propulsion, stability, and maneuverability are expected to trade off during swimming^13,38,39^. Fins that do not generate direct thrust, such as the dorsal fin and flippers, likely increase drag, whereas stability provided by fixed fins such as the dorsal fin may come at the expense of maneuverability. Consistent with this trade-off, cetacean flipper morphology has been linked to habitat and feeding behavior. In some odontocetes, relatively large flippers likely enhance maneuverability and reflect adaptation to riverine environments^13,16^. In baleen whales, by contrast, interspecific variation in flipper area is thought to reflect differences in feeding behavior arising from a trade-off between propulsion and maneuverability^14^. However, these links have largely been inferred from fin shape and area alone, with little assessment of whole-body hydrodynamic performance. Assessing whether particular dorsal fin– flipper combinations bias species toward different stability regimes is therefore essential for linking fin morphology to ecology. Such patterns must also be evaluated in a phylogenetic framework, because apparent ecological associations may reflect shared ancestry rather than repeated functional evolution.

Here, we investigated whether morphological diversity in cetacean dorsal fins, together with variation in flipper morphology and posture, generates interspecific differences in whole-body stabilization, and whether these differences are associated with ecological adaptation and evolutionary history. First, using computational fluid dynamics (CFD) simulations under different flipper postures, we assessed the whole-body stabilizing effects of the dorsal fin in five representative cetacean species spanning major differences in dorsal fin area and position, body size, and ecology. By quantifying the yawing and rolling moments generated by each body component under small yaw disturbances, we identified the dorsal fin and flipper variables underlying static postural stability. We then used geometrically approximated comparative variables to test whether these traits were associated with ecotypes defined by habitat and feeding behavior across cetaceans. Finally, we interpreted these patterns in a phylogenetic framework to infer the evolutionary origins of stabilization mechanisms associated with dorsal fin morphological diversity.

## Results

### Body component effects on yaw and roll stability

CFD simulations in ANSYS Fluent quantified drag, yawing and rolling moments under slight yaw disturbances across different flipper postures in five cetacean species, as well as in hypothetical killer whale models with modified dorsal fin area and position (Fig. 1A). Fig. 1B shows the drag coefficient and normalized yawing and rolling moments of each body component (dorsal fin, flippers, caudal fin, and trunk) under folded and 50-degree anhedral extended flipper postures in a steady 1 m s⁻¹ flow at a 5° yaw disturbance angle, with the center of volume of each model used as the center of rotation. Flipper extension increased whole-body drag in all species, with the largest increase in the killer whale model (6.0%). At 0° yaw angle, moments were negligible because the models remained bilaterally symmetrical (Supplementary Fig. SR1). At 5° and 10° yaw angles, yawing and rolling moments increased approximately proportionally across components without reversing direction, and subsequent analyses therefore focused on the 5° yaw angle condition.

**Figure 1.**
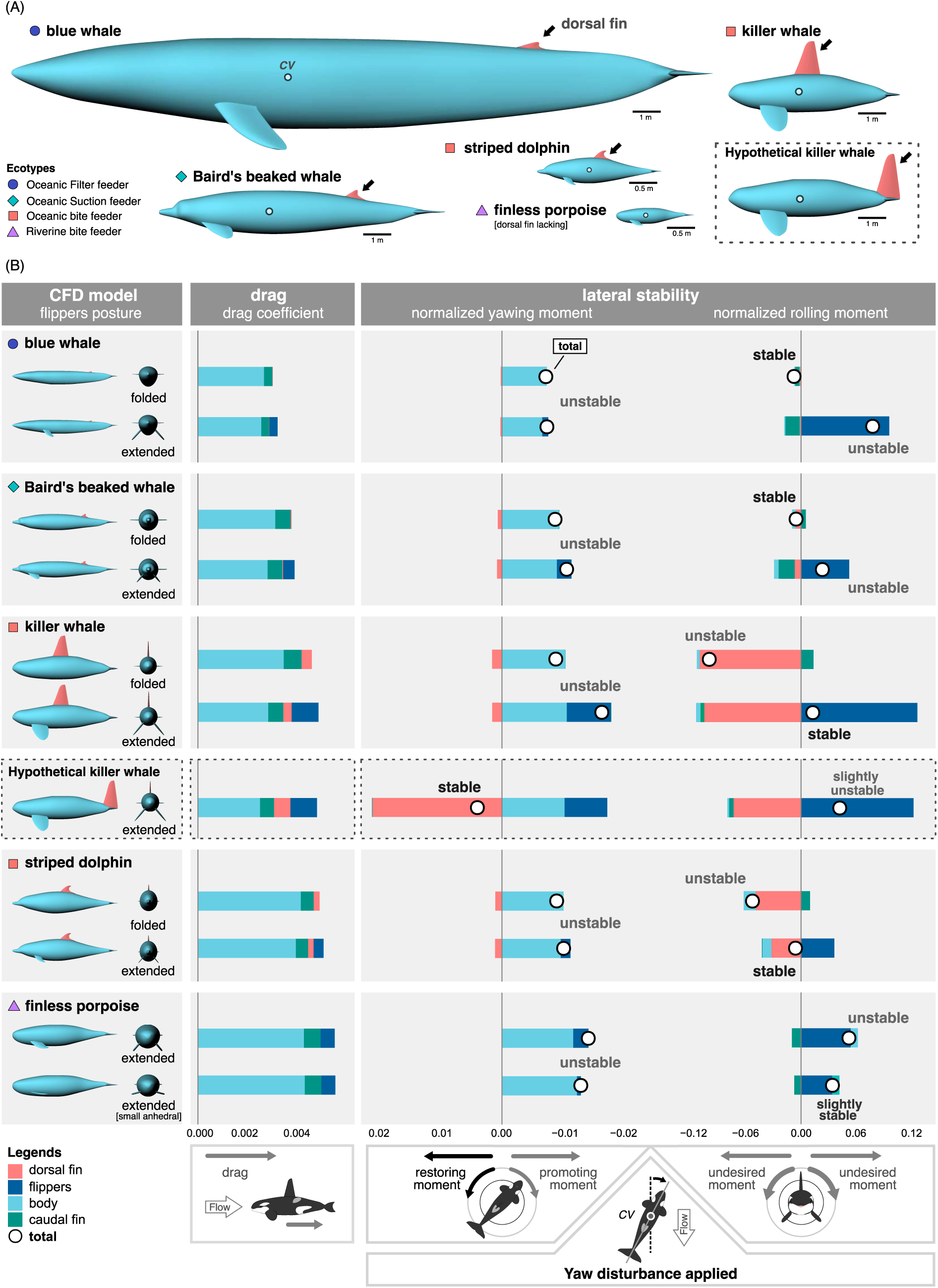
CFD simulation results for five cetacean species and a hypothetical killer whale with manipulated dorsal fin area and position. (A) CFD models of the five species and the hypothetical killer whale are shown, with the center of volume (*CV*) overlaid on each model. Plots indicate ecotypes defined by habitat and feeding behavior, and arrows indicate the dorsal fins. The finless porpoise lacks a dorsal fin. (B) Drag coefficients and normalized yawing and rolling moments about the center of rotation, approximated by the *CV*, were calculated by CFD for each body component under a 5° yaw disturbance across different flipper postures. Flipper extension increased drag relative to the folded posture. Yaw destabilization was driven primarily by the trunk regardless of flipper posture, whereas the dorsal fin was too small and too close to the center of rotation to generate sufficient restoring torque. Consequently, the whole-body normalized total yawing moment remained destabilizing in all real models (normalized total yawing moment [*M*_yaw_] > 0). Restoring yaw moments occ,wvf/..urred only in the hypothetical killer whale, in which the dorsal fin was shifted to the most posterior position and enlarged to at least 1.25 times its original area (*M*_yaw_ < 0). In contrast, rolling moments depended on the balance between dorsal fin and flipper contributions. With the flippers folded close to the trunk, species with small dorsal fins, such as the blue whale and Baird’s beaked whale, showed negligible total rolling moments (normalized total rolling moment [*M*_roll_] ≈ 0), whereas species with large dorsal fins, including the killer whale and striped dolphin, exhibited dorsal fin-induced rolling moments. When the flippers were extended at a large anhedral angle (50-degree), flipper-generated moments counterbalanced these rolling moments in species with large dorsal fins, bringing *M*_roll_ close to zero; in species with small dorsal fins, however, flipper extension instead generated undesired rolling moments. In the finless porpoise, extension at a small anhedral angle (10-degree) reduced the rolling moment requirement by approximately 60% relative to the 50-degree posture, bringing *M*_roll_ closer to zero.

For yawing moments, all models except the finless porpoise, which lacks a dorsal fin, exhibited restoring moments generated by the dorsal fin, whereas the trunk and flippers generated promoting moments (Fig. 1B, normalized yawing moment). The contribution of the caudal fin was negligible. In all species, the trunk generated the largest promoting moment, and the restoring moment from the dorsal fin was insufficient to offset it, resulting in a net promoting normalized total moment in all real models under the modeled static disturbance (normalized total yawing moment [ *M*_yaw_ ] > 0; uppercase *M* indicates normalized moment). Even in the killer whale, which exhibited the largest dorsal fin–derived restoring moment, its magnitude was only 14.9% of the trunk-derived promoting moment. Because the flippers were located anterior to the center of rotation, flipper extension further increased the promoting yawing moment relative to the folded-flipper posture. In the hypothetical killer whale model, a net restoring total moment occurred only when the dorsal fin was shifted to the most posterior position and enlarged to more than 1.25 times the area of the real model (Supplementary Fig. SR2).

Rolling moments were generated primarily by the dorsal fin and flippers, and their combined magnitude varied markedly with flipper posture (Fig. 1B, normalized rolling moment). The contributions of the trunk and caudal fin were small. Even in the striped dolphin, which exhibited the largest trunk effect, the trunk-generated moment was only 27.2% of that generated by the extended flippers. In the folded-flipper posture, species with very small dorsal fins, such as the blue whale (normalized dorsal fin area, *A*_*D*_ = 0.003; uppercase *A* indicates normalized area) and Baird’s beaked whale (*A*_*D*_ = 0.010), generated only weak rolling moments, with total moment values close to zero. By contrast, the killer whale (*A*_*D*_ = 0.10) and striped dolphin (*A*_*D*_ = 0.041) exhibited much larger rolling moments, attributable to their large dorsal fins (normalized total rolling moment [*M*_roll_] = −0.09 and −0.03, respectively). In the extended-flippers posture with a 50-degree anhedral angle, the blue whale (normalized dorsal fin to flippers area ratio [*DFR*] = 0.037), Baird’s beaked whale (*DFR* = 0.15), and finless porpoise (lacking a dorsal fin) had flippers substantially larger than the dorsal fin. As a result, the flipper-generated moments dominated and produced large net rolling moments (*M*_roll_ = 0.07, 0.03, and 0.03, respectively). By contrast, in the killer whale (*DFR* = 0.53) and striped dolphin (*DFR* = 0.87), the smaller disparity between dorsal fin and flipper areas led to near cancellation between dorsal-fin and flipper moments, bringing the total moment close to zero (*M*_roll_ = −0.004 and −0.006, respectively). This near-zero balance persisted across increasing anhedral angles up to 90 degrees (Supplementary Fig. SR3).

The finless porpoise lacks a dorsal fin and swims with its flippers extending at a small anhedral angle (∼10 degrees)^36^. In the 10-degree model, the flipper-generated rolling moment decreased to 62.7% of that of the 50-degree model, reducing the total rolling moment by about half and shifting it closer to zero. Conversely, in the killer whale, reducing the flipper anhedral angle similarly weakened the flipper-generated moment, preventing it from balancing the large dorsal-fin moment and thereby eliminating the near-zero balance observed at angles above 50 degrees (Supplementary Fig. SR3). The rolling moments in the hypothetical killer whale model with altered dorsal fin position were less pronounced than the yawing moments. Nevertheless, they varied with the dorsal trunk height relative to the body axis, and under an equal-area dorsal fin, the rolling moment was maximized when the dorsal fin was positioned as in the actual specimen (Supplementary Fig. SR2).

### Ecotype-level variation in dorsal fin and flippers proportions

The CFD simulations indicated that dorsal fin area and position had little influence on static yaw stability, whereas the dorsal fin affected roll stability in concert with the flippers; particularly under the extended-flipper posture, roll stability was determined by the area ratio between the dorsal fin and the flippers (*DFR*). To examine whether these functional relationships extend across cetaceans and are associated with ecological divergence, we compared normalized dorsal fin area (*A*\*_*D*_), normalized flipper area (*A*\**_F_*), and their area ratio (*DFR*^∗^) among 81 extant species assigned to five ecotypes based on habitat and feeding behavior. These variables were estimated from a database of external morphometric measurements of stranded cetaceans using simplified geometric approximations, with asterisks indicating approximated values. Although these variables tended to be underestimated relative to the direct measurements reflecting biomechanical responses ( *A*_*D*_ , *A*_*F*_ , and *DFR* ), the discrepancies were small compared with the magnitude of fin morphological diversity across cetaceans and did not substantially alter the relative size relationship between the dorsal fin and the flippers (Supplementary Fig. SC1).

For yaw stability, the hypothetical killer whale model showed that the yawing moment generated by the dorsal fin scaled with both normalized dorsal fin area (*A*_*D*_) and the normalized distance from the center of rotation to the dorsal fin tip (*DP*_*C*_; Supplementary Fig. SR2). Analogous to the vertical tail volume coefficient used in aircraft^40,41^, we defined the product of these two variables as the Fin Moment Index (*FMI*), a morphological index of static yaw-restoring capacity:

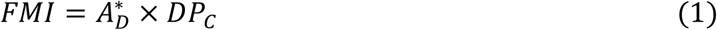

where *A*_D_* is the geometrically approximated normalized dorsal fin area. Across extant cetaceans, *FMI* was consistently much lower than in the hypothetical yaw-stable killer whale model and in the vertical tails of two representative aircraft, reflecting the generally small size of the dorsal fin and its proximity to the center of rotation (Fig. 2A). No extant species possessed dorsal fin morphology within the range required to generate a net restoring yaw moment, indicating that static yaw-restoring capacity was fundamentally constrained across cetaceans despite variation among ecotypes.

**Figure 2.**
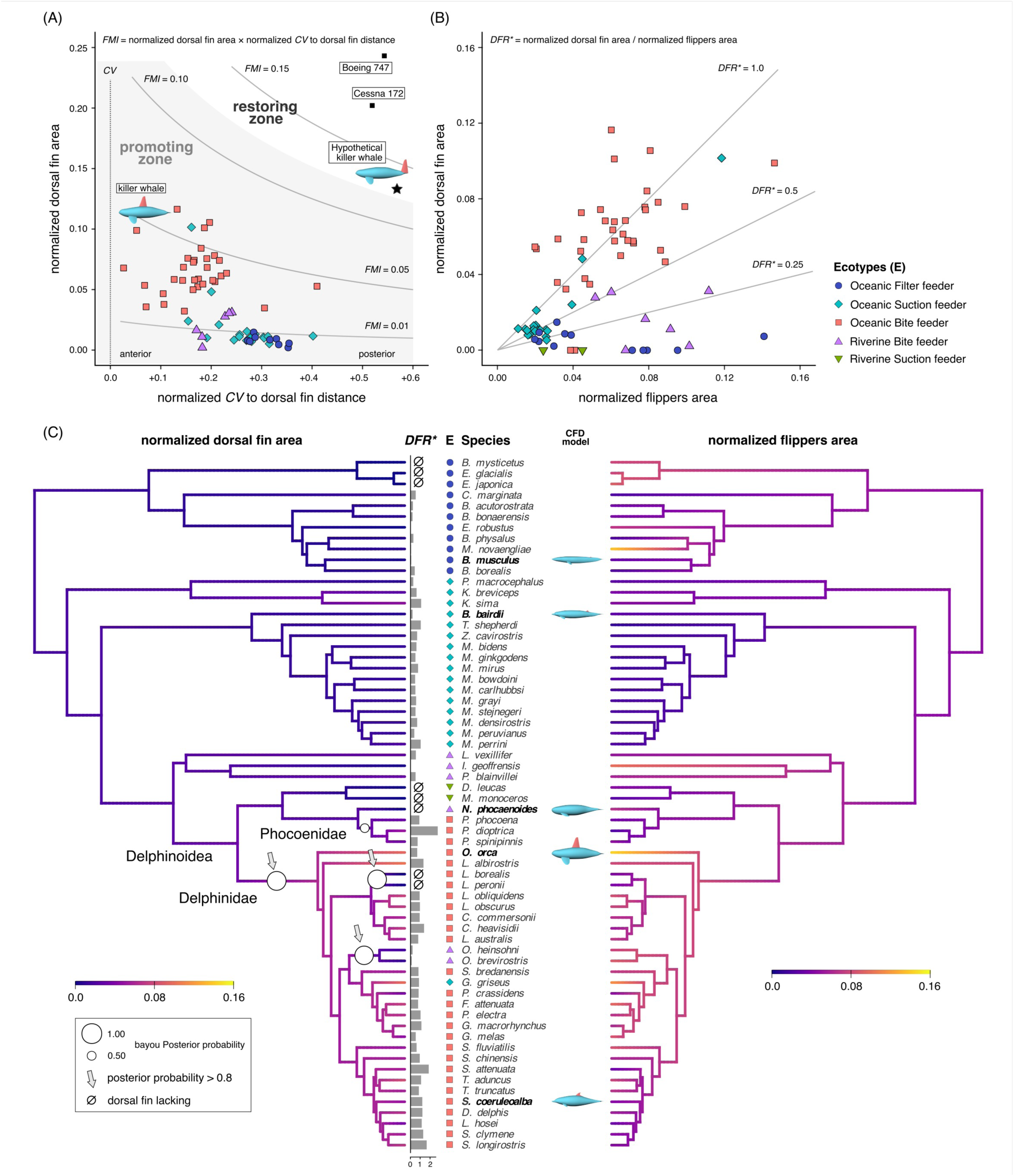
Ecotype- and phylogeny-based comparison of dorsal fin and flippers morphology related to postural stability across cetaceans. (A) Distribution of the Fin Moment Index (*FMI* = normalized dorsal fin area [*A*_D_*] × normalized distance from the center of rotation to dorsal fin [*DP*_*C*_ ]) across cetacean ecotypes, with an aircraft vertical stabilizer shown for comparison. The center of rotation was assumed to be located at 0.4 of body length from the tip of the snout. Yaw-stabilizing potential increases with *FMI*. Based on CFD results for a hypothetical killer whale model that achieved yaw stability, regions were classified as either a restoring zone or a promoting zone. No extant cetacean reached the restoring zone comparable to that of the hypothetical killer whale or aircraft. (B) The relationship between *A*_D_* and normalized flippers area (*A*_F_*) across ecotypes. Rolling moments under yaw disturbance depend on the projected area ratio between the dorsal fin and flippers (*DFR*^∗^). Combined with the CFD results, this ratio predicts ecotype-specific rolling behavior. Oceanic filter feeders possess much smaller dorsal fins relative to flippers, implying strong rolling when the flippers are extended at large anhedral angles. In contrast, oceanic bite feeders have similar dorsal fin and flipper areas (*DFR*^∗^ ≈ 1), allowing their moments to offset each other. (C) Ancestral reconstruction of *A*_D_* and *A*_F_* under the best-fitting OU model based on the molecular phylogeny of McGowen et al. (2020) [M20], with Bayou-inferred evolutionary shifts shown as circles on branches. Circle size indicates posterior probability, and only shifts with posterior probabilities ≥ 0.4 are shown. Bayou detected three shifts in *A*_D_* (arrows) with posterior probabilities > 0.8: an enlargement near the base of Delphinidae, bringing *A*_D_* close to *A*_F_* (*DFR*^∗^ ≈ 1), followed by reductions in *Lissodelphis* (*L. borealis* and *L. peronii*) and *Orcaella* (*O. brevirostris* and *O. heinsohni*). This enlargement suggests a lineage-specific acquisition of a dorsal fin large enough to counteract rolling moments generated by extended flippers at large anhedral angles. No shifts with posterior probabilities > 0.8 were detected for *A*_F_*.

In contrast, variation in dorsal fin area produced clear ecotype-level differences in potential roll stability, as quantified by *DFR*^∗^(Fig. 2B, Pairwise test results in Supplementary Table S3). Oceanic bite feeders had relatively large dorsal fins and significantly higher *DFR*^∗^ than the other ecotypes, with most individuals clustering around *DFR*^∗^ ≈ 1, indicating broadly comparable dorsal fin and flipper areas. Oceanic suction feeders had relatively small dorsal fins and flippers; their *DFR*^∗^was significantly lower than that of oceanic bite feeders but significantly higher than that of oceanic filter feeders. Oceanic filter feeders and riverine bite feeders showed a similar pattern, with dorsal fins markedly smaller than the flippers and no significant difference between these ecotypes. Although statistical comparisons were not performed for riverine suction feeders because of the limited sample size, their relative fin areas and *DFR*^∗^ followed a trend similar to that of riverine bite feeders.

Several taxa deviated from this broad association between *DFR*^∗^and ecotype (Supplementary Fig. SC2). The right whale dolphins (*Lissodelphis* sp.), classified as oceanic bite feeders, lack a dorsal fin and therefore exhibited very low *DFR*^∗^. In contrast, Risso’s dolphin (*Grampus griseus*), an oceanic suction feeder, possessed a relatively large dorsal fin comparable to those of oceanic bite feeders, resulting in high *DFR*^∗^ . Dall’s porpoise (*Phocoenoides dalli*) (*DFR*^∗^ = 2.75) and the spectacled porpoise (*Phocoena dioptrica*) (*DFR*^∗^ = 2.60), both classified as oceanic bite feeders, also had dorsal fins markedly larger than their flippers. These exceptions indicate that ecotype captures broad trends in fin proportions, but does not fully account for species-level variation.

### Phylogenetic distribution and evolutionary shifts in fin metrics

To examine the evolutionary distribution of the comparative fin metrics identified above, we estimated evolutionary models and reconstructed ancestral states for dorsal fin presence, *A*_D_*, *A*_F_* , and ecotype using two molecular phylogenies from McGowen et al.^42,43^, hereafter referred to as M09 and M20, respectively. In addition, phylogenetic shifts in *A*_D_* and *A*_F_* were inferred using bayou and OUwie analyses.

Among 71 extant cetacean species with resolved phylogenetic positions, dorsal fin loss was observed in nine species (12.7%), including members of Balaenidae, gray whale (*Eschrichtius robustus*), Monodontidae, finless porpoise, and right whale dolphins. For dorsal fin acquisition and loss, the equal-rate (ER) model provided the best fit for both phylogenies (Supplementary Table S4). Ancestral state reconstruction using the phytools estimated a >99% probability that the common ancestor of extant cetaceans already possessed a dorsal fin, indicating that dorsal fin loss occurred independently in multiple lineages (Supplementary Fig. SC3).

The normalized areas of dorsal fins and flippers showed distinct phylogenetic shifts (Fig. 2C). For 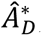, the Ornstein–Uhlenbeck (OU) model best fit both molecular phylogenies of McGowen et al. (2009, 2020) (Supplementary Table S5). Bayou and OUwie supported a two-step OU model with three selective regimes^44^. Bayou detected three evolutionary shifts with posterior probabilities >0.8, corresponding to an increase in *A*_D_* in Delphinidae and a subsequent decrease toward the clade including snubfin dolphins *Orcaella* sp. and right whale dolphins (Supplementary Figs. SC4 and SC7). OUwie identified the OUM model assuming distinct optima per regime as the best supported model (Supplementary Table S6). The estimated half-life was short relative to the tree height (0.06 for M20; 0.14 for M09), indicating a rapid convergence to regime-specific optima. In the OU model, the common ancestor of extant cetaceans had *A*_D_* in ∼1.7% of the trunk projected area, comparable to extant Ziphiidae. These analyses indicate that, although dorsal fin presence is broadly shared among extant cetaceans, marked dorsal fin enlargement evolved selectively within only a subset of Delphinidae.

For *A*_F_* , no phylogenetic shift as clear as that observed for *A*_D_* was detected. Under M20, Bayou did not identify distinct selective regimes (Supplementary Fig. SC5). Under M09, by contrast, Bayou identified four independent shifts with posterior probabilities exceeding 0.8, corresponding to Balaenidae, the humpback whale (*Megaptera novaeangliae*), the Ganges River dolphin (*Platanista gangetica*), and the killer whale. However, these shifts were confined to terminal branches, and OUwie estimated a relatively long half-life (0.3 of tree height), indicating weaker and less coordinated change than in *A*_D_* (Supplementary Fig. SC8). In the OU model, the common ancestor of extant cetaceans was estimated to have an *A*_F_* of ∼7.3% of trunk projected area, comparable to that of extant Delphinidae. These results indicate that evolutionary change in dorsal fin area was more strongly structured by phylogeny than was change in flipper area.

For ecotypes, the ER model was selected in both phylogenies (Supplementary Table S4), and no particular ecotype was reconstructed for the common ancestor of extant cetaceans. River-adapted lineages arose repeatedly across the phylogeny, whereas posterior probabilities for other ecotypes increased sharply along specific branches (Supplementary Fig. SC10). Oceanic bite feeding was reconstructed within Delphinoidea, with the highest support appearing by at least the base of Delphinidae (M20: 76.1% at Delphinoidea and >99% at Delphinidae; M09: 93.8% at Delphinoidea and >99% at Delphinidae). Combined with the *A*_D_* results, this pattern indicates that dorsal fin enlargement linked to roll stability evolved in only a subset of lineages within the oceanic bite-feeding niche.

## Discussion

CFD simulations of whole-body models with varied dorsal fin shapes and flipper postures showed that the dorsal fin does not function as a universal static stabilizer in cetaceans, under yaw-disturbance conditions. Furthermore, the stabilization contribution is primarily expressed in roll rather than in yaw. Although the cross-sectional shapes and aspect ratios of dorsal fins and flippers vary across species, previous studies have reported only modest differences in vertical force and drag coefficients^27,29,30^. Because hydrodynamic forces and the resulting moments are directly proportional to these coefficients, the overall trends observed in our simulations are unlikely to change substantially even with higher model resolution or greater analytical precision.

Under yaw disturbances, yawing moments were dominated by the trunk in all models, whereas restoring moments, which were consistently too small to counteract the destabilizing effect of the trunk, were generated by the dorsal fin regardless of fin area or position. The simulations included species expected to maximize *FMI*, such as the killer whale, which has the largest dorsal fin area among extant cetaceans, and the blue whale, whose dorsal fin is positioned most posteriorly. Even in these cases, dorsal fin–mediated restoring moments remained small. Comparative morphological analyses likewise showed that all extant cetaceans occupy a low-*FMI* region of morphospace, consistent with the limited yaw-restoring potential of the dorsal fin. Thus, the finding that the dorsal fin contributes little to static yaw stability is robust unless the assumed center of rotation is displaced to an extreme position.

By contrast, roll effects varied strongly with the balance between dorsal fin- and flipper-generated moments. When the flippers were folded to reduce drag, species with small dorsal fins, such as the blue whale and Baird’s beaked whale, generated only negligible rolling moments. Species with large dorsal fins, such as the killer whale and striped dolphin, by contrast, showed large dorsal fin-induced rolling moments. Extending the flippers at a large anhedral angle increased drag in all species but produced divergent outcomes. In species with large dorsal fins, flipper-generated moments counterbalanced dorsal fin-induced rolling moments, whereas in species with small dorsal fins, the same extension generated undesired rolling moments. These results indicate that dorsal fin function cannot be understood in isolation and that its principal hydrodynamic effect, particularly on roll stability, depends on mechanical coupling with flipper morphology and posture.

Taken together, the CFD and comparative analyses suggest three roll-stabilization strategies in cetaceans (Fig. 3): folded-flippers, dorsal fin, and small anhedral flipper strategies. The folded-flippers strategy minimizes moment generation by keeping the flippers closely folded against the trunk with a small, hydrodynamically minor dorsal fin. The dorsal fin strategy counterbalances large flipper-generated moments with a dorsal fin of roughly comparable projected area when the flippers are extended at a large anhedral angle. The small anhedral flipper strategy reduces flipper-generated rolling moments by extending the flippers more laterally at a small anhedral angle while retaining a small or absent dorsal fin. Notably, both the folded-flippers and small anhedral flipper strategies can reduce the total rolling moments through the flipper posture alone, without relying on a large dorsal fin. The folded-flippers strategy is likely to be the most efficient during straight-line swimming, because it minimizes protrusions that increase whole-body drag^16,45^ and reduces the muscular effort required to hold the flippers extended^46^. By contrast, the dorsal-fin strategy likely incurs higher hydrodynamic and postural costs because it requires both a relatively large dorsal fin and sustained flipper extension. The small-anhedral-flipper strategy avoids reliance on a large dorsal fin and may therefore reduce drag, but it appears to provide only limited mitigation of rolling moments.

**Figure 3.**
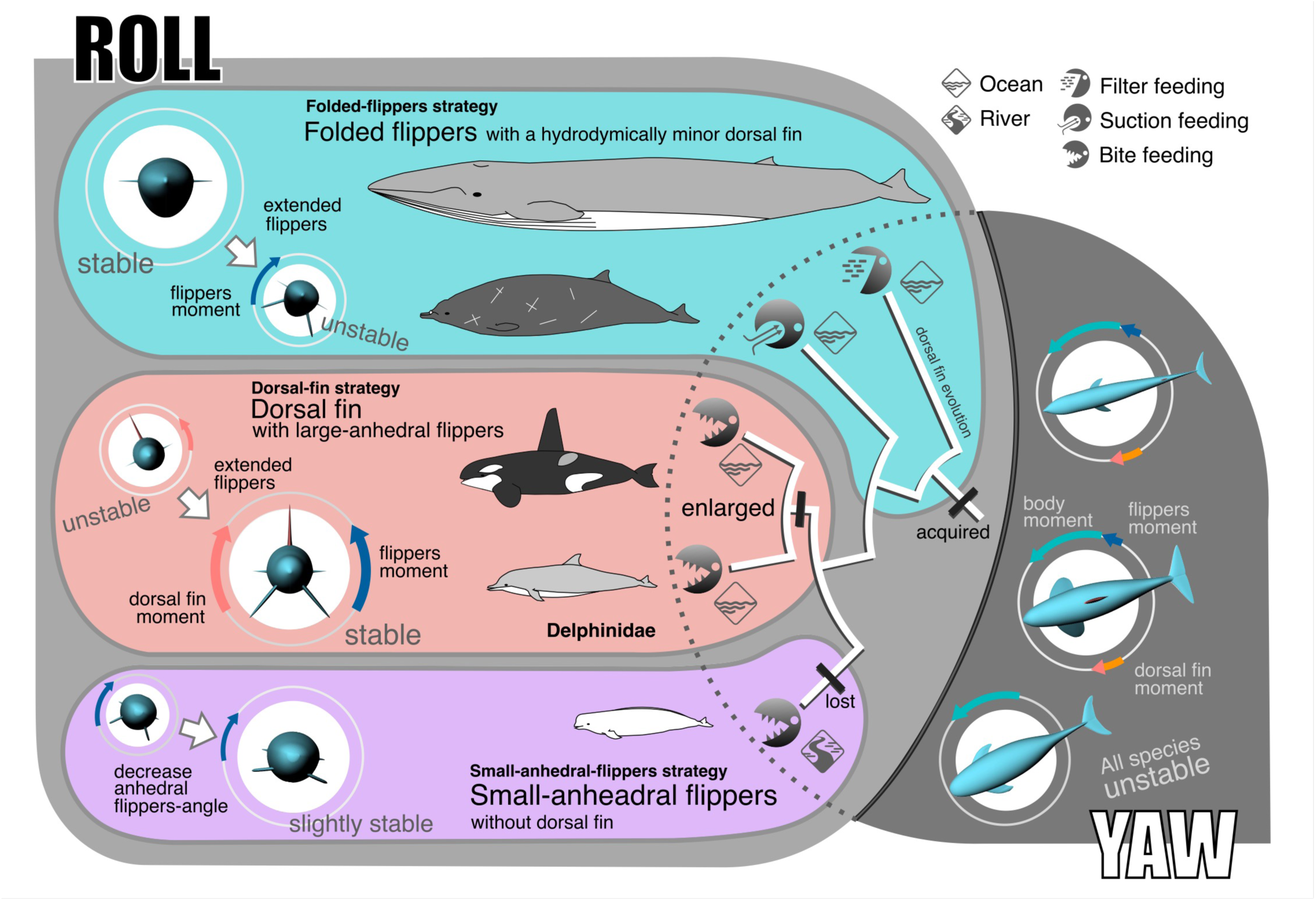
Dorsal fin in flipper-coupled roll stabilization strategies of cetaceans. Cetacean dorsal fins contribute little to static yaw stability, but are associated with three flipper-coupled strategies of roll stabilization linked to ecological adaptation. Filter- and suction-feeding species maintain roll stability with small dorsal fins and folded flippers, a configuration that also reduces drag during straight-line swimming (folded-flipper strategy). Oceanic bite feeders, by contrast, achieve both roll stability and the maneuverability required for precise prey tracking through a large dorsal fin and persistently extended flippers (dorsal fin strategy). Riverine species reduce roll instability with a small dorsal fin and flippers extended at a small anhedral angle, while avoiding strong fixed rolling moments from the dorsal fin and retaining greater maneuverability than in the dorsal fin strategy (small anhedral flipper strategy). In both the folded-flipper and small anhedral flipper strategies, roll stability can be maintained primarily by flipper posture alone, even without a dorsal fin. Phylogenetic comparisons further indicate that the common ancestor of extant cetaceans already possessed a dorsal fin, but enlargement to an area comparable to that of the flippers occurred only in a limited number of lineages, particularly within Delphinidae. Together, these findings suggest that the dorsal fin did not function as a universal stabilizer throughout cetacean evolution. Instead, lineage-specific evolutionary enlargement of the dorsal fin may have enabled persistent flipper extension, thereby facilitating the emergence of a dorsal fin–mediated stabilization strategy and the exploitation of novel oceanic foraging niches.

These strategies can be interpreted as alternative resolutions of a trade-off between minimizing drag and roll moments during routine swimming and retaining flipper-mediated control authority for maneuvering. The dorsal fin strategy, which combines a large dorsal fin with sustained flipper extension, is mechanically costly but may enhance instantaneous maneuverability through greater flipper-mediated control^7,13^. Many oceanic bite-feeding species exhibit relatively large dorsal fins and only small differences between dorsal fin and flipper area. Because this ecotype requires precise prey pursuit, it combines high-speed swimming with agile maneuvering^47,48^. Although extended flippers facilitate rapid angle-of-attack adjustments during turning, maintaining them in an extended posture can increase rolling moments during straight-line swimming^33,34,49^. A dorsal fin with an area comparable to the projected flipper area is therefore consistent with balancing roll during routine swimming while preserving maneuvering capacity. This configuration, however, also increases drag and likely requires greater mechanical power than morphologies with smaller or absent dorsal fins.

Oceanic filter-feeding species, oceanic suction feeders, and many river-adapted taxa generally possess dorsal fins much smaller than their flippers. In filter- and suction-feeders, which likely require less routine maneuvering than oceanic bite feeders^13^, this pattern is most consistent with a folded-flippers strategy during straight swimming, in which small dorsal fins and folded flippers reduce drag and rolling moments^36,50^. In baleen whales, the relatively large flippers may instead be extended only transiently during feeding or specific maneuvers, when large rolling moments are advantageous^16,34,51^. Suction-feeding taxa, particularly beaked whales, also show small dorsal fins and flippers, together with low *DFR*^∗^, consistent with specialization for drag reduction rather than for balancing moments between the dorsal fin and flippers. Socket-like depressions at the flipper base further suggest a limited contribution of the flippers to stability^25^. River-adapted species, including the finless porpoise and beluga, show small anhedral flipper angles, and active flipper-assisted turning has been observed in the Amazon River dolphin^52^. Adaptation to shallow, structurally complex riverine environments may favor a small-anhedral-flippers strategy, in which dorsal fin reduction suppresses excessive roll-correcting moments while extended flippers enhance maneuverability^13^. The relatively limited reduction in rolling moment compared with the other two strategies may indicate that strong stability is less important in these taxa and that maneuverability is prioritized.

Comparison with phylogenetic data suggested that the common ancestor of extant cetaceans already possessed a dorsal fin. However, dorsal fin–mediated roll stabilization appears to have evolved only in specific lineages, likely within Delphinidae, under selective pressures favoring rapid dorsal fin enlargement. This marked enlargement may in turn have enabled persistent flipper extension, thereby facilitating the adoption of a dorsal fin–mediated stabilization strategy in association with oceanic bite-feeding niches. Some Phocoenidae also possess relatively large dorsal fins associated with a similar ecotype, leaving unresolved whether this strategy evolved independently in Delphinidae and Phocoenidae or originated at the base of Delphinoidea, which includes Delphinidae, Phocoenidae, and Monodontidae. In any case, the phylogenetic analyses indicate that roll stabilization mediated by the dorsal fin was not a universal feature of extant cetaceans, but rather a derived function restricted to particular lineages after the origin of the dorsal fin itself. Convergent secondary reductions in dorsal fin area within Delphinidae further suggest that evolutionary flexibility in dorsal fin morphology facilitated diversification in roll-stabilization strategies.

This lineage-specific pattern also suggests that the functions of the cetacean dorsal fin cannot be reduced to a single universal stabilizing role. Dorsal fins have also been proposed to contribute to thermoregulation via internal vasculature^53–55^, visual display associated with sexual dimorphism^56,57^, and vortex generation that may enhance propulsive efficiency^58,59^. These functions may have contributed to the origin and diversification of the dorsal fin before lineage-specific enlargement made its role in roll stabilization more prominent in some taxa. Our findings suggest that the cetacean dorsal fin is not the product of a single evolutionary pathway dedicated to hydrodynamic stabilization, but rather a multifunctional structure shaped by the sequential addition, transformation, and lineage-specific emphasis of multiple functions.

### Limitations

This study has several limitations. In particular, the dorsal fin- and flipper-mediated roll-stabilization strategies proposed here do not fully capture the complexity of cetacean swimming and should therefore be interpreted as broad functional trends associated with ecotypes. First, our analyses evaluated static stability using rigid-body models under steady flow and did not incorporate full swimming kinematics. Accordingly, the CFD results indicate only that the dorsal fin made a limited contribution to static yaw restoration under the modeled conditions, rather than that yaw stability is generally unimportant in vivo. Flippers may contribute to yaw stabilization through adjustable angles of attack produced by internal and external rotation at the shoulder joint. Tail flexibility and torsion may also contribute to yaw control^60–62^. Future studies should therefore evaluate dynamic stability using unsteady analyses that incorporate tail and flipper motion, as well as trunk flexibility, through fluid–structure interaction.

Second, behavioral data on flipper posture, particularly anhedral angle, remain scarce for many species. In this study, we used 50° as a standardized comparative posture based on observations in small cetaceans^35^, although this may not represent the habitual posture of all taxa. Our results suggest that the posture required to balance dorsal fin- and flippers-derived rolling moments depends on the relative proportions of these structures. In species in which flipper area exceeds dorsal fin area, modulation of flipper posture may help balance rolling moments between the two. By contrast, in taxa such as the spectacled porpoise, Dall’s porpoise, and humpback dolphins (*Sousa* sp.), in which dorsal fin area clearly exceeds flipper area, adjustment of flipper posture alone may be insufficient to offset dorsal fin-derived rolling moments. This, in turn, suggests that such species may rely on species-specific strategies for postural roll stabilization. A deeper understanding of the hydrodynamic function of the dorsal fin in coordination with the flippers will therefore require species-level biologging and kinematic data that jointly quantify flipper posture and ecological context^63^.

Third, no cetacean fossils clearly preserving dorsal fins have been reported, precluding direct assessment of dorsal fin evolution in extinct taxa. We therefore focused on extant species and used living diversity as a comparative framework. However, because extinct lineages underwent lineage-specific radiations and extinctions through evolutionary time^64,65^, phylogenetic patterns derived from extant species alone should not be interpreted as a complete history of dorsal fin enlargement. Enlarged dorsal fins may have been more widespread in extinct clades than extant diversity suggests. Future advances in fossil soft-tissue reconstruction and evolutionary developmental biology may allow more reliable inference of dorsal fin structure and developmental origins in extinct taxa. In addition, the relationship identified here between dorsal fin area and flipper posture may provide a basis for indirectly inferring dorsal fin function in fossil species from preserved shoulder morphology.

Despite these limitations, our results demonstrate that, although the cetacean dorsal fin is a homologous structure across species, variation in its area can produce markedly different stabilizing effects. They further show that its function cannot be understood from fin morphology alone, but must be evaluated from a whole-body perspective that explicitly considers its mechanical relationship with the flippers. This framework may also be extended to other aquatic vertebrates bearing multiple fins, especially clades with both fixed and movable fins, such as ichthyosaurs and sharks^8,15,10^. More broadly, these findings underscore the difficulty of explaining the origin and diversification of fin structures in aquatic vertebrates solely in hydrodynamic terms and highlight the complexity of their fin evolution.

In conclusion, this study challenges the long-standing view that the cetacean dorsal fin functions as a universal vertical stabilizer and shows that dorsal fin stabilization, in coupling with the flippers, is restricted to particular phylogenetic lineages and ecological contexts. The dorsal fin contributes little to whole-body yaw stability, and its role in roll stabilization appears to be largely confined to Delphinidae, in which rapid evolutionary enlargement of the dorsal fin, coupled with flipper extension, may have facilitated the emergence of a novel stabilization strategy associated with oceanic bite feeding. In most other lineages, by contrast, roll stability is maintained during swimming by flipper posture alone, without requiring the dorsal fin. These findings suggest that the evolutionary history of the dorsal fin cannot be explained by stabilization alone, but is better understood as part of a multifunctional evolutionary pathway.

## Methods

### 3D modeling

CFD simulations were conducted on whole-body three-dimensional (3D) models of five cetacean species selected to span the diversity of dorsal fin morphology, body size, ecology, and phylogenetic position in extant cetaceans: blue whale (*Balaenoptera musculus*; family: Balaenopteridae; *BL*: 24.80 m; ecotype: oceanic filter feeder), Baird’s beaked whale (*Berardius bairdii*; Ziphiidae; 10.20 m; oceanic suction feeder), killer whale (*Orcinus orca*; Delphinidae; 6.05 m; oceanic bite feeder), striped dolphin (*Stenella coeruleoalba*; Delphinidae; 2.21 m; oceanic bite feeder), and finless porpoise (*Neophocaena asiaeorientalis*; Phocoenidae; 1.39 m; riverine bite feeder). This set included the killer whale, with one of the largest dorsal fins among extant cetaceans; the blue whale, with one of the smallest and most posteriorly positioned dorsal fins among extant cetaceans; and the finless porpoise, which lacks a dorsal fin entirely. Flipper posture was varied from folded and extended at anhedral angles ranging from 0 to 90 degrees. Additional hypothetical killer whale models with modified dorsal fin area and position were also simulated.

Three-dimensional models were constructed from image data and external morphometric measurements using Rhinoceros 8 (Robert McNeel & Associates) and its integrated visual programming environment, Grasshopper (Supplementary Fig. SM1). Each model consisted of four separately constructed components: the trunk, paired flippers, dorsal fin, and caudal fin. These were assembled into a complete whole-body model. All models were assumed to be bilaterally symmetrical, rigid, and in a straight swimming posture. The body axis was defined as the line from the tip of the snout to the caudal fin notch, and this distance was used as body length (*BL*). A right-handed coordinate system was adopted, with the positive X-axis directed from head to tail, the positive Y-axis from the body midline toward the right side, and the positive Z-axis dorsally.

Because the body axis in cetaceans does not necessarily pass through the center of each trunk cross section in the transverse plane, the trunk was reconstructed by separately generating dorsal and ventral components from half-elliptical cross sections defined along the body axis and fitted to the corresponding outlines (Supplementary Fig. SM1A). Cross-sectional radii were determined as the perpendicular distances from the body axis to NURBS curves fitted to dorsal and ventral outlines extracted from image data. To reduce distortion caused by body curvature in photographs, the body axis and outlines were geometrically straightened before reconstruction. A provisional body-axis NURBS curve defined from anatomical landmarks was resampled, straightened along the X-axis while preserving local distances, and used to remap the measured perpendicular distances to generate outlines corresponding to a straight body axis. The resulting dorsal and ventral outlines were then used to reconstruct the trunk.

The fins were modeled separately from the trunk (Supplementary Fig. SM1B–D). Their leading and trailing edges were traced from photographs and reference images with NURBS curves, and hydrofoil sections approximating cetacean fins were extruded along these outlines^27^: the E836 profile for the flippers and dorsal fin, and the S1048 profile for the caudal fin. All fins were fixed at an angle of attack of 0° under the 0° yaw condition. Among fins, only the flippers were allowed to vary in posture: the anhedral angle was changed among simulation cases by rotation about the shoulder joint, whereas the dorsal and caudal fins were kept immobile (Supplementary Fig. SM1E).

The sources of image data and morphometric measurements used for model construction differed among species (Supplementary Fig. SM2). For the striped dolphin (KPM-NFM 8005) and finless porpoise (KPM-NFM 8003), models were based on photographs and external measurements of single freshly stranded individuals. For the killer whale, Baird’s beaked whale, and blue whale, individual components were reconstructed from multiple specimens and then scaled and positioned using external morphometric measurements.

Two flipper postures were defined. In the folded posture, the flippers were omitted from the exposed surface to represent an idealized condition in which they were closely appressed to the trunk. In the extended posture, each flipper was rotated anhedrally about the shoulder joint while maintaining constant area and the minimum sweep angle, defined as the configuration in which the elbow contacted the trunk surface^15,35,37,52^. Sensitivity analyses for the killer whale and blue whale included a folded model and extended models with anhedral angles from 0-degree to 90-degree at 20-degree intervals (Supplementary Fig. SR3). Because flipper-generated moments reached a plateau above the approximately 50° anhedral posture reported for small odontocetes^35,37^, interspecific comparisons were conducted using folded models and extended models with a 50-degree anhedral posture (Fig. 1B). For the finless porpoise, an additional model with a 10-degree anhedral posture was included based on observational records.

Additional hypothetical models were constructed for the killer whale to examine the effects of dorsal fin position and size. In addition to the real killer whale model, hypothetical models were generated in which the normalized distance from the center of rotation to the dorsal fin tip ( *DP*_*C*_ ; defined below) varied from −0.21 to +0.54 at intervals of approximately 0.10 while dorsal fin area was held constant. In the most posterior condition (*DP*_*C*_ = +0.54), dorsal fin area was further increased to 1.25 and 2.00 times the original value by extending the fin dorsally while keeping basal length constant (Supplementary Fig. SR2). In all hypothetical models, flipper posture was fixed at a 50° anhedral extended position. For the finless porpoise, an additional model was constructed to evaluate the hydrodynamic effect of the dorsal ridge. Ridge height and width were set to twice the values measured from a stranded specimen, and the ridge was generated by extruding a semi-elliptical vertical cross section along the dorsal trunk surface.

Based on these models, morphological variables relevant to the yaw-disturbance analyses were defined. The normalized distance from the center of rotation to dorsal fin tip (*DP*_*C*_ ) was defined along the body axis as the distance from the center of rotation to the dorsal fin tip, normalized by *BL*:

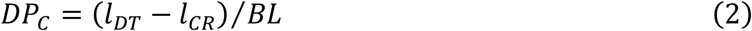

where *l*_*DT*_ and *l*_*CR*_ are the distances from the snout tip to the dorsal fin tip and to the center of rotation, respectively. Because the exact center of rotation is difficult to determine, the center of volume of each model was used as a proxy. In cetaceans, the centers of mass and buoyancy are thought to be nearly coincident, and the center of mass of the bottlenose dolphin (*Tursiops* sp.) has been reported at approximately 0.41*BL* from the snout tip^13,68^. Consistent with this, the centers of volume of the five CFD models were located at 0.38–0.42 *BL*s from the snout tip. In addition, normalized fin areas were defined relative to the lateral projected area of the trunk (*a*_*T*,*lat*_) because this study focuses on yaw disturbances. The normalized areas of the dorsal fin (*A*_*D*_) and flipper (*A*_*F*_) were calculated as the lateral projected area of each fin divided by *a*_*T*,lat_. For paired flippers, *A*_*F*_ was defined as the sum of the lateral projected areas of both sides:

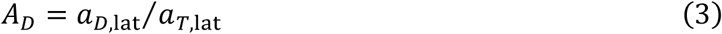

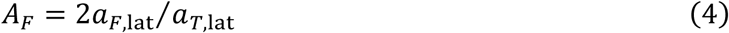

For the caudal fin, which extends along the left–right axis, *A*_*C*_ was defined as the dorsal projected area divided by the lateral projected area of the body:

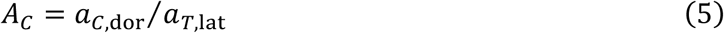

The dorsal fin to flippers area ratio (*DFR*) was then calculated as:

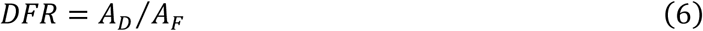

where *a*_*T*,lat_, *a*_*D*,lat_, *a*_*F*,lat_, and *a*_*C*,dor_ denote the lateral projected areas of the trunk, dorsal fin, flipper, and the dorsal projected area of the caudal fin, respectively. All measurements were performed in Rhinoceros (Supplementary Fig. SC1), and the resulting values for each model are presented in Supplementary Table S1.

### CFD simulations

CFD simulations were conducted using ANSYS 2023 R2 (Ansys Inc.). The fluid domain was generated in ANSYS DesignModeler by subtracting the cetacean model from the surrounding fluid region using a Boolean operation. Each model was placed in a sufficiently large cuboidal computational domain (280 m × 120 m × 140 m in the X-, Y-, and Z-directions, respectively) to minimize boundary effects (Supplementary Fig. SM3). Boundary conditions comprised a velocity inlet at the upstream face, a pressure outlet at the downstream face, symmetry conditions on the top and bottom surfaces, and a no-slip wall on the model surface. At the lateral boundaries in the Y-direction, the upstream side was defined as a velocity inlet and the opposite side as a pressure outlet. Meshes were generated in ANSYS Meshing primarily with tetrahedral elements. A boundary-layer mesh consisting of 20 prism layers was applied to the model surface, with the first-layer thickness adjusted to maintain y⁺ < 5. To reduce the total mesh count while maintaining high resolution near the body, the Body of Influence function was used with up to four nested cuboidal refinement regions surrounding the model. Depending on the model, the total number of elements ranged from approximately 7 to 17 million.

Flow simulations were performed in ANSYS Fluent. Pressure–velocity coupling was handled using the SIMPLE algorithm, and turbulence was modeled by solving the Reynolds-averaged Navier– Stokes equations with the SST *k*–*ω* model. Seawater was used as the working fluid, with a density (*ρ*) of 1026 kg m⁻³ and a dynamic viscosity (*μ*) of 0.00117 kg m⁻¹ s⁻¹, and the inlet velocity (*U*) was set to 1.0 m s⁻¹. Reynolds number was defined as *Re* = *ρUBL*/*μ*, where *ρ* is fluid density, *U* is flow velocity, *μ* is dynamic viscosity, and *BL* is body length. At this inlet velocity, *Re* ranged from 1.2 × 10^6^ (finless porpoise) to 21.8 × 10^6^ (blue whale) across the models. Each simulation was run for at least 4,000 iterations and terminated once the fluid forces acting on each body component in the X, Y, and Z directions had stabilized, as indicated by a stable plateau and a root mean square (RMS) fluctuation below 10⁻³. RMS values were calculated over a moving window of 50 consecutive iterations after excluding the first 500 iterations, and final force values were obtained by averaging the last 100 iterations. Inspection of the whole-body pressure distributions confirmed that no localized anomalous values attributable to mesh discontinuities or other obvious numerical artifacts were present in any model. Under yaw disturbance, pressure differences were observed across the two sides of the dorsal fin and flippers in association with local flow-velocity changes, consistent with the generation of lift and the resulting moments (Supplementary Fig. SM4).

The center of volume of each model was adopted as the center of rotation in this study. Moments about the model body axes, *X*_*B*_ , *Y*_*B*_ , and *Z*_*B*_ , were defined as roll, pitch, and yaw, respectively (Supplementary Fig. SM5). Because body posture was fixed throughout the simulations, these body axes coincided with the spatial axes (*X*, *Y*, and *Z*). Yaw disturbances were reproduced by varying the *X*- and *Y*-components of the inflow velocity, and simulations were conducted at yaw angles of 0°, 5°, and 10°. Hydrodynamic forces acting on each body component, namely the trunk, dorsal fin, flippers, and caudal fin, as well as the moments generated by those forces, were calculated along the three spatial axes. For each component *i* (*i* = dorsal fin, flippers, caudal fin, and trunk), these hydrodynamic forces and moments were nondimensionalized as a drag coefficient (*C*_*D*,*i*_ ) and normalized rolling, pitching, and yawing moments (*M*_roll,*i*_, *M*_pitch,*i*_, *M*_yaw,*i*_).

The drag coefficient of component *i* was defined as:

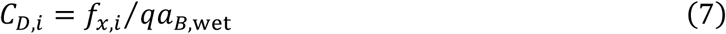

where *f*_*x*,*i*_ is the force acting on component *i* in the *X*-direction, *a*_*B*,wet_ is the wetted surface area of the whole body, and *q* is the dynamic pressure, defined as *q* = 1⁄2*ρU*^2^, where *ρ* is fluid density and *U* is inflow velocity. The normalized rolling moment of component *i* was defined as:

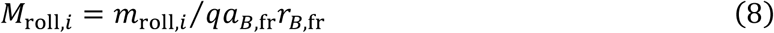

where *m*_roll,*i*_ is the rolling moment generated by component *i* about the *X*_*B*_-axis, *a*_*B*,fr_ is the frontal projected area of the whole body, and *r*_*B*,fr_ is the maximum distance from the body axis on the frontal whole-body projection. The normalized pitching moment of component *i* was defined as:

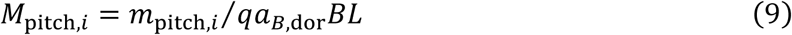

where *m*_pitch,*i*_ is the pitching moment generated by component *i* about the *Y*_*B*_-axis, and *a*_*B*,dor_ is the dorsal projected area of the whole body. The normalized yawing moment of component *i* was defined as:

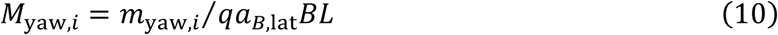

where *m*_yaw,*i*_ is the yawing moment generated by component *i* about the *Z*_*B*_-axis, *a*_*B*,lat_ is the lateral projected area of the whole body, and *BL* is body length. The quantities *a*_*B*,lat_ , *a*_*B*,fr_ , *a*_*B*,dor_ , and *r*_*B*,fr_ were calculated for each model using Rhinoceros (Supplementary Fig. SC1).

Because all component moments were normalized using the same whole-body reference quantities, the total moment of the whole body was defined as the sum of the normalized component moments:

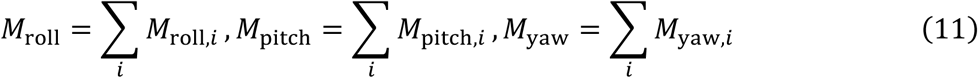

Thus, *M*_roll_ , *M*_pitch_ , and *M*_yaw_ represent the whole-body normalized rolling, pitching and yawing moments, respectively.

The numerical approach was validated separately for drag and moment calculations. For drag validation, two prolate spheroid models (4154 and 4159) reported by Gertler^69^ were simulated under the same numerical conditions as the cetacean models. To match *Re* range of the cetacean models, the prolate spheroid models were geometrically scaled to total lengths of 2.74, 5.48, 13.7, and 21.9 m (*Re* = 2.4 × 10^6^, 4.8 × 10^6^, 12.0 × 10^6^, and 19.2 × 10^6^, respectively); for these models, *BL* was defined as the maximum length along the *X*_*B*_-axis. The computed drag values were within 5% of the corresponding experimental measurements, and the cetacean CFD models at comparable Reynolds numbers showed similar drag values (Supplementary Fig. SM6). Moment calculations were validated by confirming near-zero values under the symmetric 0° yaw-angle condition.

In addition, mesh sensitivity was evaluated using the killer whale model as a representative case with a prominent dorsal fin and strong fin-induced moments. Three mesh conditions, comprising approximately 3.1, 8.7, and 14.6 million elements, were compared. Because the differences in rotational moments among these conditions were within 1%, the intermediate mesh, with approximately 7 million elements, was adopted for the main analyses (Supplementary Fig. SM7). The effect of swimming speed was likewise assessed by conducting additional simulations at inlet velocities of 0.5, 1.0 and 2.0 m s⁻¹ (*Re* = 2.7 × 10^6^, 5.3 × 10^6^, 10.6 × 10^6^). Although increasing flow velocity increased the absolute magnitudes of the moments, it did not substantially alter the overall pattern of normalized moments (Supplementary Fig. SM8). Finally, the effect of the dorsal ridge in the finless porpoise was smaller than that of the dorsal fins in the other models and had little influence on the conclusions of this study (Supplementary Fig. SM9).

### Morphometric data compilation and normalization

External morphometric data including fin lengths, from 6,580 adult individuals of 81 cetacean species (14 families) were compiled from published sources and standardized to 0.1 cm (the subset used in this study is provided in Supplementary Table S2, and the full data source is available in DELPHI). Measurement points followed the standardized methods for smaller cetaceans^66^, and sources without explicit citation to this protocol were also included when the landmarks were consistent. When multiple source-specific mean values were available for a species, an overall species mean was calculated as a whole sample size mean. Although dorsal fin morphology exhibits sexual and geographic variation^67^, dorsal fin presence is not sex-specific and was not distinguished in this study. The distance from the center of rotation to the dorsal fin tip (*DP*_*C*_ ) was calculated in the same manner as for the CFD models (Eq. 2), with the center of rotation fixed at 0.40*BL* from the snout tip for all species. This value is consistent with both the CFD models (0.38–0.42*BL*) and the reported center of mass of the bottlenose dolphin (0.41*BL*)^13^. Normalized fin areas, including those of the dorsal fin (*A*\**_D_*), flippers (*A*\**_F_*), and caudal fin (*A*\**_C_*), were calculated from species mean values using geometric approximations based on two linear measurements (Supplementary Fig. SC1). The lateral projected area of the trunk (*a*_T,lat_*) was approximated as an ellipse defined by *BL* and maximum body (trunk) height (*BH*). Because *BH* is not included in the standardized methods^66^, a circular cross-section was assumed in this study, with family-specific fineness ratios (*FR = BL/BW*, where *BW ≈ BH*) from Fish and Rohr^16^ used to estimate *BH*; values from closely related families were substituted when necessary. Dorsal fin and caudal fin areas were approximated as triangles. The lateral projected area of the dorsal fin (*a*_D,lat_*) was calculated as half the product of its maximum height and base length, and the dorsal projected area of the caudal fin (*a*_C,dotr_*) as half the product of maximum span and base length. Flipper lateral area was approximated as a rhombus with a 50-degree anhedral posture. Accordingly, the lateral projected area of the left flipper (*a*\**_f,left,lat_*) was calculated as half the product of the leading-edge length and maximum width of the left flipper, multiplied by sin 50^∘^; total flipper area was defined as twice this value, assuming bilateral symmetry. Normalized fin areas were then calculated as follows:

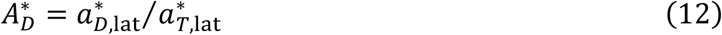

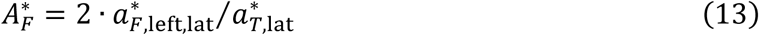

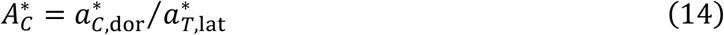

Species lacking a dorsal fin were assigned *A*\**_D_* = 0.0001 to allow continuous trait analysis, below the minimum observed in cetaceans (0.002 in blue whale). The dorsal fin to flippers area ratio (*DFR*^∗^) was calculated as:

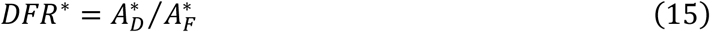

For the two representative aircraft used for comparison, *FMI* was derived from image data for the Cessna 172^70^ and Boeing 747^71^ by quantifying the projected body area, the relative area of the vertical tail, and its distance from the center of rotation.

### Fin morphology across ecotypes

Extant cetacean species were classified into ecotypes based on habitat and feeding behavior (Table S2). Habitats were categorized as oceanic or riverine. Species documented by Jefferson et al.^72^ and Würsig et al.^73^ as residents of or regularly entering river systems were assigned to the riverine category. Feeding behavior was classified as filter, suction, or bite feeding by cross-referencing multiple studies that primarily relied on behavioral observations^74–77^. These ecotypes were used as broad comparative categories rather than as exhaustive representations of all within-species behavioral variation. Although six theoretical combinations of habitat and feeding behavior exist (2 × 3), no extant species exhibited a riverine filter feeding ecotype.

Inter-ecotype differences in *A*\**_D_*, *A*\**_F_*, *DFR*^∗^ , and *A*\**_C_* were assessed using the Steel–Dwass multiple comparison procedure implemented in the R package PMCMRplus; pairwise differences were considered significant at *p* < 0.05 (Supplementary Table S3). The riverine suction feeding ecotype was excluded from statistical comparison due to insufficient sample size (n = 2). No significant differences among ecotypes were detected for *A*\**_C_*.

### Ancestral reconstruction and evolutionary shifts

The presence of dorsal fin, *A*\**_D_*, *A*\**_F_*, *A*\**_C_*, and ecotypes was analyzed within a phylogenetic framework of extant cetaceans (M20^43^, M09^42^) to estimate evolutionary models and ancestral states. Dorsal fin presence (coded as finned or finless) and ecotype were treated as discrete traits, whereas *A*\**_D_* , *A*\**_F_*, and *DFR*^∗^ were treated as continuous traits. Analyses were restricted to the 71 extant species with resolved phylogenetic positions in these trees. To examine the evolutionary relationships between the dorsal fin and flippers areas, species lacking the relevant continuous variables were excluded from the corresponding comparisons.

For discrete traits, evolutionary models were fitted using the fitMk function in the R package phytools under three transition-rate models: equal-rates (ER), symmetric-rates (SYM), and all-rates-different (ARD). Model fit was evaluated using Akaike information criterion (AIC), and the model with the lowest AIC was selected as the best-fitting model. Ancestral states and branch-specific transition probabilities were then estimated by performing 10,000 stochastic character-mapping simulations with the make.simmap function in phytools (Supplementary Figs. SC3 and SC10).

For continuous traits, the model fit was assessed using the fitContinuous function in the R package geiger under the Brownian motion (BM), OU, and early burst models. When the OU model was selected as the best-fitting model (lowest AIC), the following analyses were conducted. Bayesian MCMC analyses were performed using the R package bayou based on the OU model to identify the branches associated with potential adaptive regime shifts^44^. For the prior distributions, multiple independent MCMC chains with different priors were run, and the optimal parameters were selected based on the marginal likelihood (Supplementary Figs. SC4–SC9). For each trait, five independent chains of 1,000,000 generations were executed, with the first 20% discarded as burn-in. Convergence was assessed using the Gelman-Rubin diagnostic (*R*^ < 1.1), and branches with posterior probabilities exceeding 0.8 were identified as adaptive regimes. For *A*_C_*, although the OU model was selected as the best-fitting model, no branches with posterior probabilities exceeding 0.8 were detected by Bayou in either phylogeny (Supplementary Figs. SC6 and SC9).

To evaluate whether the adaptive regimes estimated by bayou correspond to rapid evolutionary shifts, additional model comparisons were conducted using the R package OUwie. Phylogenies were partitioned according to the regime configurations inferred by bayou using the paintSubTree function, and OUM (multiple *θ*), OU1 (single *θ*), and BM models were fitted. When the OUM model was selected as the best-fitting model (lowest AIC), the estimated half-life parameters were compared with the total tree height to assess the rate of evolutionary shifts in a phylogenetic context.

## Supporting information

Supplemental Informations

## Acknowledgements

We sincerely thank N. Toyoda, Y. Goto, D.M. Kikuchi, H. Kajiyama, S. Fujiwara, and the members of the Ecology Group at Nagoya University for their helpful comments and support. We also thank N. Mitsuoka and A.R. Smith for their assistance with stranded cetacean surveys. We thank OKA Design Studio for assistance with graphic design. This work was supported by JSPS KAKENHI (JP23KJ1113 and JP21H05294), Anri Scholarship, and Toyoaki Scholarship Foundation. We gratefully acknowledge Ansys, Inc. for providing ANSYS software through the Ansys Academic Research Partnership program.

## Conflicts of Interest

The authors declare no conflicts of interest.

## Data Availability Statement

Supplementary figures and tables are provided in the Supplementary Information. The main 3D models used for the CFD simulations are uploaded at Sketchfab (https://skfb.ly/pIAZo). The external morphometric measurement data and their corresponding source references are available as DELPHI at figshare (https://doi.org/10.6084/m9.figshare.31998957). The CFD simulation outputs and the R code used for analysis and visualization are also deposited at figshare (https://doi.org/10.6084/m9.figshare.31999257). All data underlying the figures and tables are available from these repositories.

## Author Contributions

T.O. conceived the study, designed the research, performed the CFD simulations and comparative analyses, visualized the data, secured funding, and wrote the original draft. M.M. contributed to study design, data interpretation, and manuscript revision. F.N. contributed to data acquisition for 3D model construction, interpretation of the results, and manuscript revision. K.Y. supervised the project, secured funding, and revised the manuscript. All authors discussed the results and approved the final version of the manuscript.

## Notes

https://skfb.ly/pIAZo

https://doi.org/10.6084/m9.figshare.31998957

https://doi.org/10.6084/m9.figshare.31999257

### Competing Interest Statement

The authors have declared no competing interest.

