## Supplemental Informations for "Dorsal fins are not universal stabilizers in cetaceans: limited yaw effects and flipper-coupled roll stability"

**Contents**

Symbol list **2**

Tables (Table S1–S6) **3**

Figures

CFD methods (Figures SM1–SM9) **7**

CFD results (Figures SR1–SR3) **16**

Ecotype and phylogenetic comparisons (Figures SC1–S10) **19**

### List of symbols

**Body dimensions and positions**

$BL$ body length

$BH$ body (trunk) maximum height

$l_{DT}$ the distances from the snout tip to the dorsal fin tip

$l_{CR}$ the distances from the snout tip to the center of rotation of the body

$CV$ the center of volume

$DP_C$ normalized distance from the center of rotation (using $CV$ as a proxy) to dorsal fin tip

**Projected indices**

$a_{T,\mathrm{lat}}$ lateral projected trunk area

$a_{D,\mathrm{lat}}$ lateral projected dorsal fin area

$a_{F,\mathrm{left,lat}}$ lateral projected area of the left flipper

$a_{C,\mathrm{dor}}$ dorsal projected caudal fin area

$A_D$ normalized dorsal fin area

$A_F$ normalized flippers area

$A_C$ normalized caudal fin area

$DFR$ normalized dorsal fin to flippers area ratio $[A_D/A_F]$

**Geometrically approximated projected indices**

$a^*_{T,\mathrm{lat}}$ geometrically approximated lateral projected trunk area

$a^*_{D,\mathrm{lat}}$ geometrically approximated lateral projected dorsal fin area

$a^*_{F,\mathrm{left,lat}}$ geometrically approximated lateral projected area of the left flipper

$a^*_{C,\mathrm{dor}}$ geometrically approximated dorsal projected caudal fin area

$A^*_D$ geometrically approximated normalized dorsal fin area

$A^*_F$ geometrically approximated normalized flippers area

$A^*_C$ geometrically approximated normalized caudal fin area

$DFR^*$ geometrically approximated normalized dorsal fin to flippers area ratio $[A^*_D/A^*_F]$

$FMI$ Fin moment index $[A^*_D \times DP_C]$

**Whole-body reference quantities**

$a_{B,\mathrm{wet}}$ wetted surface area of the whole-body

$a_{B,\mathrm{lat}}$ whole-body lateral projected area

$a_{B,\mathrm{fr}}$ whole-body frontal projected area

$a_{B,\mathrm{dor}}$ whole-body dorsal projected area

$r_{B,\mathrm{fr}}$ maximum distance from the body axis on the frontal whole-body projection

**Flow quantities**

$Re$ Reynolds number

$q$ dynamic pressure

$\rho$ density [1026 kg m$^{-3}$]

$\mu$ dynamic viscosity [0.00117 kg m$^{-1}$ s$^{-1}$]

$U$ inlet fluid velocity

**Component-wise forces and moments**

$i$ body component index (dorsal fin, flippers, caudal fin and trunk)

$f_{x,i}$ force on component $i$ in the $X$-direction

$C_{D,i}$ drag coefficient of component $i$

$m_{\mathrm{roll},i}$ rolling moment of component $i$

$m_{\mathrm{pitch},i}$ pitching moment of component $i$

$m_{\mathrm{yaw},i}$ yawing moment of component $i$

$M_{\mathrm{roll},i}$ normalized rolling moment of component $i$

$M_{\mathrm{pitch},i}$ normalized pitching moment of component $i$

$M_{\mathrm{yaw},i}$ normalized yawing moment of component $i$

$M_{\mathrm{roll}}$ whole-body normalized total rolling moment

$M_{\mathrm{pitch}}$ whole-body normalized total pitching moment

$M_{yaw}$ whole-body normalized total yawing moment

**Table S1** Morphological variables of the 3D cetacean and prolate spheroid models used in the CFD simulations. The table includes Species name, model name, FP [flipper posture; folded or extended at an anhedral angle], $BL$ [body length (m)]; $P_C$, normalized position of the center of volume from the snout tip; $DP_C$, normalized distance from the center of volume to the dorsal fin tip; $k_D$, dorsal fin size scaling factor; $a_{D,\mathrm{lat}}$, lateral projected dorsal fin area (m$^2$); $a_{F,\mathrm{lat}}$, lateral projected area of the left flipper (m$^2$); $a_{C,\mathrm{dor}}$, dorsal projected caudal fin area (m$^2$); $A_D$, normalized dorsal fin area (dimensionless); $A_F$, normalized flipper area (dimensionless); $A_C$, normalized caudal fin area (dimensionless); $a_{B,\mathrm{wet}}$, wetted surface area of the whole body (m$^2$); $a_{B,\mathrm{lat}}$, lateral projected area of the whole body (m$^2$); $a_{B,\mathrm{dor}}$, dorsal projected area of the whole body (m$^2$); $a_{B,\mathrm{fr}}$, frontal projected area of the whole body (m$^2$); and $r_{B,\mathrm{fr}}$, maximum distance from the body axis on the frontal whole-body projection (m). Positive and negative $DP_C$ values indicate that the dorsal fin tip is located posterior and anterior to the center of rotation, respectively. ∅ indicates values not applicable because of fin absence. For the sunameri $F_{50}$Ridge model, the listed variables correspond to the non-homologous dorsal ridge structure.

| Species | model name | FP | $BL$ | $P_C$ | $DP_C$ | $k_D$ | $a_{D,\mathrm{lat}}$ | $a_{F,\mathrm{left,lat}}$ | $a_{C,\mathrm{dor}}$ | $A_D$ | $A_F$ | $A_C$ | $a_{B,\mathrm{wet}}$ | $a_{B,\mathrm{lat}}$ | $a_{B,\mathrm{dor}}$ | $a_{B,\mathrm{fr}}$ | $r_{B,\mathrm{fr}}$ |
|---|---|---|---|---|---|---|---|---|---|---|---|---|---|---|---|---|---|
| Blue whale | whale Folded | folded | 24.8 | 0.41 | +0.36 | 1.00 | 0.16 | ∅ | 2.10 | 0.0028 | — | 0.036 | 177 | 58.3 | 51.5 | 7.86 | 2.61 |
| | whale $F_0$ | 0-degree | 24.8 | 0.41 | +0.36 | 1.00 | 0.16 | 2.92 | 2.10 | 0.0028 | 0 | 0.036 | 189 | 58.3 | 56.0 | 8.87 | 3.51 |
| | whale $F_{10}$ | 10-degree | 24.8 | 0.41 | +0.36 | 1.00 | 0.16 | 2.92 | 2.10 | 0.0028 | 0.017 | 0.036 | 189 | 58.3 | 55.9 | 8.87 | 3.62 |
| | whale $F_{30}$ | 30-degree | 24.8 | 0.41 | +0.36 | 1.00 | 0.16 | 2.92 | 2.10 | 0.0028 | 0.050 | 0.036 | 189 | 58.8 | 55.0 | 8.87 | 3.76 |
| | whale $F_{50}$ | 50-degree | 24.8 | 0.41 | +0.36 | 1.00 | 0.16 | 2.92 | 2.10 | 0.0028 | 0.075 | 0.035 | 189 | 59.3 | 54.2 | 8.87 | 3.78 |
| | whale $F_{70}$ | 70-degree | 24.8 | 0.41 | +0.36 | 1.00 | 0.16 | 2.92 | 2.10 | 0.0027 | 0.092 | 0.035 | 189 | 59.8 | 52.5 | 8.87 | 3.70 |
| | whale $F_{90}$ | 90-degree | 24.8 | 0.41 | +0.36 | 1.00 | 0.16 | 2.92 | 2.10 | 0.0027 | 0.098 | 0.035 | 189 | 59.9 | 51.5 | 8.87 | 3.51 |
| Baird’s beaked whale | beaked-whale Folded | folded | 10.2 | 0.42 | +0.30 | 1.00 | 0.11 | ∅ | 0.71 | 0.010 | ∅ | 0.064 | 36.7 | 11.1 | 10.8 | 2.02 | 1.41 |
| | beaked-whale $F_{50}$ | 50-degree | 10.2 | 0.42 | +0.30 | 1.00 | 0.11 | 0.50 | 0.71 | 0.0099 | 0.067 | 0.063 | 38.5 | 11.3 | 11.0 | 2.12 | 1.41 |
| Killer whale | killer-whale Folded | folded | 6.05 | 0.42 | +0.08 | 1.00 | 0.68 | ∅ | 0.44 | 0.11 | ∅ | 0.070 | 19.7 | 6.27 | 5.68 | 1.48 | 1.89 |
| | killer-whale $F_0$ | 0-degree | 6.05 | 0.42 | +0.08 | 1.00 | 0.68 | 0.86 | 0.44 | 0.11 | 0 | 0.070 | 23.1 | 6.27 | 7.03 | 1.67 | 1.89 |
| | killer-whale $F_{10}$ | 10-degree | 6.05 | 0.42 | +0.08 | 1.00 | 0.68 | 0.86 | 0.44 | 0.11 | 0.048 | 0.070 | 23.1 | 6.28 | 7.02 | 1.67 | 1.89 |
| | killer-whale $F_{30}$ | 30-degree | 6.05 | 0.42 | +0.08 | 1.00 | 0.68 | 0.86 | 0.44 | 0.10 | 0.13 | 0.067 | 23.1 | 6.54 | 6.77 | 1.67 | 1.89 |
| | killer-whale $F_{50}$ | 50-degree | 6.05 | 0.42 | +0.08 | 1.00 | 0.68 | 0.86 | 0.44 | 0.10 | 0.19 | 0.065 | 23.1 | 6.78 | 6.41 | 1.67 | 1.89 |
| | killer-whale $F_{70}$ | 70-degree | 6.05 | 0.42 | +0.08 | 1.00 | 0.68 | 0.86 | 0.44 | 0.099 | 0.23 | 0.063 | 23.1 | 6.94 | 5.86 | 1.67 | 1.89 |
| | killer-whale $F_{90}$ | 90-degree | 6.05 | 0.42 | +0.08 | 1.00 | 0.68 | 0.86 | 0.44 | 0.098 | 0.25 | 0.063 | 23.1 | 7.00 | 5.67 | 1.67 | 1.89 |
| Hypothetical | killer-whale $F_{50}D_{M21}$ | 50-degree | 6.05 | 0.42 | −0.21 | 1.00 | 0.68 | 0.86 | 0.44 | 0.10 | 0.19 | 0.065 | 23.1 | 6.78 | 6.41 | 1.66 | 1.71 |
| Killer whale | killer-whale $F_{50}D_{M15}$ | 50-degree | 6.05 | 0.42 | −0.15 | 1.00 | 0.68 | 0.86 | 0.44 | 0.10 | 0.19 | 0.065 | 23.1 | 6.78 | 6.41 | 1.66 | 1.71 |
| | killer-whale $F_{50}D_{M05}$ | 50-degree | 6.05 | 0.42 | −0.05 | 1.00 | 0.68 | 0.86 | 0.44 | 0.10 | 0.19 | 0.065 | 23.1 | 6.78 | 6.41 | 1.66 | 1.71 |
| | killer-whale $F_{50}D_{15}$ | 50-degree | 6.05 | 0.42 | +0.15 | 1.00 | 0.68 | 0.86 | 0.44 | 0.10 | 0.19 | 0.065 | 23.1 | 6.78 | 6.41 | 1.67 | 1.71 |
| | killer-whale $F_{50}D_{25}$ | 50-degree | 6.05 | 0.42 | +0.25 | 1.00 | 0.68 | 0.86 | 0.44 | 0.10 | 0.19 | 0.065 | 23.1 | 6.78 | 6.41 | 1.66 | 1.71 |
| | killer-whale $F_{50}D_{35}$ | 50-degree | 6.05 | 0.42 | +0.35 | 1.00 | 0.68 | 0.86 | 0.44 | 0.10 | 0.19 | 0.065 | 23.1 | 6.78 | 6.41 | 1.65 | 1.71 |
| | killer-whale $F_{50}D_{45}$ | 50-degree | 6.05 | 0.42 | +0.45 | 1.00 | 0.68 | 0.86 | 0.44 | 0.10 | 0.19 | 0.065 | 23.1 | 6.78 | 6.41 | 1.64 | 1.71 |
| | killer-whale $F_{50}D_{55}$ | 50-degree | 6.05 | 0.42 | +0.54 | 1.00 | 0.68 | 0.86 | 0.44 | 0.10 | 0.19 | 0.065 | 23.1 | 6.78 | 6.41 | 1.61 | 1.71 |
| | killer-whale $F_{50}D_{55}A_{125}$ | 50-degree | 6.05 | 0.43 | +0.54 | 1.25 | 0.88 | 0.86 | 0.44 | 0.13 | 0.19 | 0.063 | 23.5 | 6.94 | 6.41 | 1.64 | 1.71 |
| | killer-whale $F_{50}D_{55}A_{200}$ | 50-degree | 6.05 | 0.43 | +0.54 | 2.00 | 1.10 | 0.86 | 0.44 | 0.15 | 0.19 | 0.062 | 23.7 | 7.11 | 6.41 | 1.66 | 1.84 |
| Striped dolphin | dolphin Folded | folded | 2.21 | 0.42 | +0.15 | 1.00 | 0.027 | ∅ | 0.28 | 0.041 | ∅ | 0.044 | 1.95 | 0.65 | 0.55 | 0.16 | 0.38 |
| | dolphin $F_{50}$ | 50-degree | 2.21 | 0.42 | +0.15 | 1.00 | 0.027 | 0.02 | 0.28 | 0.041 | 0.047 | 0.043 | 2.03 | 0.65 | 0.56 | 0.16 | 0.38 |
| Finless porpoise | sunameri $F_{10}$ | 10-degree | 1.39 | 0.38 | ∅ | ∅ | ∅ | 0.02 | 0.02 | ∅ | 0.022 | 0.063 | 1.04 | 0.28 | 0.32 | 0.079 | 0.26 |
| | sunameri $F_{50}$ | 50-degree | 1.39 | 0.38 | ∅ | ∅ | ∅ | 0.02 | 0.02 | ∅ | 0.095 | 0.063 | 1.04 | 0.29 | 0.30 | 0.079 | 0.26 |
| | sunameri $F_{50}$Ridge | 50-degree | 1.39 | 0.38 | ∅ | ∅ | (0.026) | 0.02 | 0.02 | (0.085) | 0.088 | 0.058 | 1.06 | 0.31 | 0.30 | 0.079 | 0.26 |
| 4159 | e4159 | — | 2.74 | — | — | — | — | — | — | — | — | — | 1.79 | 0.057 | — | — | — |
| 4159 (2 times) | e4159-2 | — | 5.48 | — | — | — | — | — | — | — | — | — | 7.16 | 0.23 | — | — | — |
| 4159 (5 times) | e4159-5 | — | 13.7 | — | — | — | — | — | — | — | — | — | 44.8 | 1.41 | — | — | — |
| 4159 (8 times) | e4159-8 | — | 21.9 | — | — | — | — | — | — | — | — | — | 114.6 | 3.62 | — | — | — |
| 4154 | e4154 | — | 2.74 | — | — | — | — | — | — | — | — | — | 4.65 | 0.36 | — | — | — |
| 4154 (2 times) | e4154-2 | — | 5.48 | — | — | — | — | — | — | — | — | — | 18.6 | 1.45 | — | — | — |
| 4154 (5 times) | e4154-5 | — | 13.7 | — | — | — | — | — | — | — | — | — | 116.2 | 9.04 | — | — | — |
| 4154 (8 times) | e4154-8 | — | 21.9 | — | — | — | — | — | — | — | — | — | 297.7 | 23.2 | — | — | — |

**Table S2** Measurement-based comparative variables for extant cetacean species used in ecotype comparisons and phylogenetic comparative analyses are listed. The table includes the studied species [family and species name], sample size [N], habitat [H; ocean or river], feeding behavior [FB; filter, suction, or bite], ecotype [E; ecotype abbreviation], dorsal fin presence [finned; finned or finless], and body length [$BL$ (m)]. Variables marked with asterisks indicate geometrically approximated values: $a^*_{D,\mathrm{lat}}$, lateral projected dorsal fin area (m$^2$); $a^*_{F,\mathrm{left,lat}}$, lateral projected area of the left flipper (m$^2$); $a^*_{C,\mathrm{dor}}$, dorsal projected caudal fin area (m$^2$); $a^*_{T,\mathrm{lat}}$, lateral projected trunk area (m$^2$); $DP_C$, normalized distance from the center of rotation to the dorsal fin tip (dimensionless); $A^*_D$, normalized dorsal fin area (dimensionless); $A^*_F$, normalized flipper area (dimensionless); $A^*_C$, normalized caudal fin area (dimensionless); and $DFR^*$, normalized dorsal fin to flipper area ratio (dimensionless). Ref. (FB) indicates the reference for feeding behavior classification. ∅ indicates values not applicable because of dorsal fin absence. Positive $DP_C$ values indicate that the dorsal fin tip is positioned posterior to the center of rotation.

| family | species | N | H | FB | E | finned | $BL$ | $DP_C$ | $a^*_{D,\mathrm{lat}}$ | $a^*_{F,left,\mathrm{lat}}$ | $a^*_{C,\mathrm{dor}}$ | $a^*_{T,\mathrm{lat}}$ | $A^*_D$ | $A^*_F$ | $A^*_C$ | $DFR^*$ | Ref. (FB) |
|---|---|---|---|---|---|---|---|---|---|---|---|---|---|---|---|---|---|
| Balaenidae | *B. mysticetus* | 5 | Ocean | filter | OF | finless | 13.77 | ∅ | ∅ | 1.45 | 3.55 | 28.12 | ∅ | 0.079 | 0.126 | 0 | C1,C2 |
| | *E. glacialis* | 17 | Ocean | filter | OF | finless | 12.30 | ∅ | ∅ | 1.13 | — | 22.43 | ∅ | 0.077 | — | 0 | J,C1,C2 |
| | *E. japonica* | 12 | Ocean | filter | OF | finless | 15.63 | ∅ | ∅ | 2.25 | 3.75 | 36.22 | ∅ | 0.095 | 0.104 | 0 | J,C1,C2 |
| Neobalaenidae | *C. marginata* | 5 | Ocean | filter | OF | finned | 6.18 | +0.29 | 0.061 | 0.076 | 0.40 | 5.66 | 0.011 | 0.021 | 0.071 | 0.524 | J,C1,C2 |
| Eschrichtiidae | *E. robustus* | 12 | Ocean | filter | OF | finless | 11.99 | ∅ | ∅ | 0.84 | 1.2 | 17.95 | ∅ | 0.071 | 0.071 | 0 | J,O,C1,C2 |
| Balaenopteridae | *B. acutorostrata* | 1731 | Ocean | filter | OF | finned | 8.21 | +0.32 | 0.073 | 0.20 | 0.22 | 8.40 | 0.009 | 0.036 | 0.026 | 0.243 | O,C2 |
| | *B. bonaerensis* | 3 | Ocean | filter | OF | finned | 7.94 | +0.27 | 0.063 | 0.20 | 0.56 | 7.87 | 0.008 | 0.039 | 0.072 | 0.206 | O |
| | *B. physalus* | 681 | Ocean | filter | OF | finned | 20.01 | +0.35 | 0.29 | 0.65 | 2.26 | 49.93 | 0.006 | 0.020 | 0.045 | 0.29 | J,C1,C2 |
| | *M. novaengliae* | 46 | Ocean | filter | OF | finned | 12.23 | +0.28 | 0.14 | 1.71 | 2.07 | 18.63 | 0.007 | 0.141 | 0.111 | 0.052 | J,C1,C2 |
| | *B. musculus* | 722 | Ocean | filter | OF | finned | 24.80 | +0.35 | 0.17 | 1.50 | 3.59 | 76.67 | 0.002 | 0.030 | 0.047 | 0.074 | J,C |
| | *B. omurai* | 1 | Ocean | filter | OF | finned | 5.70 | +0.28 | 0.06 | 0.083 | 0.30 | 4.05 | 0.015 | 0.031 | 0.074 | 0.471 | C2 |
| | *B. borealis* | 190 | Ocean | filter | OF | finned | 15.20 | +0.33 | 0.27 | 0.42 | 1.67 | 28.80 | 0.009 | 0.022 | 0.058 | 0.426 | C2 |
| | *B. brydei* | 182 | Ocean | filter | OF | finned | 13.40 | +0.33 | 0.11 | 0.32 | 1.29 | 22.37 | 0.005 | 0.022 | 0.058 | 0.214 | C2 |
| Physeteridae | *P. macrocephalus* | 185 | Ocean | suction | OS | finned | 13.23 | +0.28 | 0.17 | 0.39 | 1.80 | 23.31 | 0.007 | 0.026 | 0.077 | 0.279 | O,C2 |
| Kogiidae | *K. breviceps* | 5 | Ocean | suction | OS | finned | 2.71 | +0.16 | 0.024 | 0.025 | — | 0.98 | 0.024 | 0.039 | — | 0.612 | O,C1,C2 |
| | *K. sima* | 13 | Ocean | suction | OS | finned | 2.31 | +0.20 | 0.034 | 0.021 | 0.064 | 0.71 | 0.048 | 0.045 | 0.090 | 1.075 | O,C2 |
| Platanistidae | *P. gangetica* | 3 | River | bite | RB | finned | 1.90 | +0.24 | 0.015 | 0.035 | — | 0.48 | 0.031 | 0.112 | — | 0.28 | O,C2 |
| Ziphiidae | *T. shepherdi* | 6 | Ocean | suction | OS | finned | 6.40 | +0.31 | 0.069 | 0.044 | 0.25 | 6.06 | 0.011 | 0.011 | 0.042 | 1.034 | J,C1,C2 |
| | *B. bairdii* | 289 | Ocean | suction | OS | finned | 10.03 | +0.32 | 0.080 | 0.25 | 1.10 | 14.91 | 0.005 | 0.026 | 0.074 | 0.206 | O,C2 |
| | *B. arnuxii* | 13 | Ocean | suction | OS | finned | 8.89 | +0.31 | — | 0.19 | — | 11.70 | — | 0.025 | — | — | C2 |
| | *B. minimus* | 3 | Ocean | suction | OS | finned | 6.71 | +0.30 | 0.072 | 0.112 | 0.49 | 6.67 | 0.011 | 0.026 | 0.074 | 0.420 | C1 |
| | *Z. cavirostris* | 26 | Ocean | suction | OS | finned | 5.56 | +0.28 | 0.048 | 0.047 | 0.30 | 4.58 | 0.010 | 0.016 | 0.065 | 0.667 | J,O,C1,C2 |
| | *I. pacificus* | 6 | Ocean | suction | OS | finned | 5.86 | +0.21 | — | — | 0.31 | 5.10 | — | — | 0.060 | — | C1,C2 |
| | *H. ampullatus* | 2 | Ocean | suction | OS | finned | 7.46 | +0.27 | 0.13 | — | — | 8.24 | 0.016 | — | — | — | C1,C2 |
| | *H. planifrons* | 1 | Ocean | suction | OS | finned | 7.45 | +0.29 | 0.11 | 0.109 | 0.69 | 8.23 | 0.013 | 0.020 | 0.083 | 0.649 | C2 |
| | *M. bidens* | 2 | Ocean | suction | OS | finned | 4.57 | +0.40 | 0.036 | 0.039 | 0.16 | 3.09 | 0.012 | 0.020 | 0.051 | 0.600 | C1,C2 |
| | *M. ginkgodens* | 6 | Ocean | suction | OS | finned | 4.76 | +0.27 | 0.027 | 0.035 | 0.22 | 3.35 | 0.008 | 0.016 | 0.066 | 0.496 | C1,C2 |
| | *M. mirus* | 3 | Ocean | suction | OS | finned | 4.89 | +0.26 | 0.040 | 0.034 | 0.18 | 3.54 | 0.011 | 0.015 | 0.050 | 0.761 | C1,C2 |
| | *M. bowdoini* | 1 | Ocean | suction | OS | finned | 4.34 | — | 0.030 | 0.041 | — | 2.79 | — | 0.011 | 0.023 | 0.471 | C1,C2 |
| | *M. carlhubbsi* | 9 | Ocean | suction | OS | finned | 5.05 | +0.25 | 0.032 | 0.042 | — | 3.78 | 0.008 | 0.017 | — | 0.492 | J,O,C1,C2 |
| | *M. layardii* | 1 | Ocean | suction | OS | finned | 5.21 | +0.30 | — | 0.048 | — | 4.02 | — | 0.018 | — | — | C1,C2 |
| | *M. hectori* | 2 | Ocean | suction | OS | finned | 3.89 | +0.22 | — | 0.029 | 0.18 | 2.24 | — | 0.020 | 0.082 | — | C1,C2 |
| | *M. densirostris* | 3 | Ocean | suction | OS | finned | 4.19 | +0.26 | 0.034 | 0.033 | 0.16 | 2.60 | 0.013 | 0.019 | 0.061 | 0.669 | C1,C2 |
| | *M. stejnegeri* | 4 | Ocean | suction | OS | finned | 4.58 | +0.31 | 0.031 | 0.041 | 0.13 | 3.10 | 0.010 | 0.020 | 0.041 | 0.504 | O,C1,C2 |
| | *M. grayi* | 53 | Ocean | suction | OS | finned | 4.74 | +0.28 | 0.036 | 0.042 | 0.28 | 3.33 | 0.011 | 0.019 | 0.083 | 0.565 | C1,C2 |
| | *M. perrini* | 1 | Ocean | suction | OS | finned | 3.90 | +0.22 | 0.048 | 0.03 | 0.17 | 2.25 | 0.021 | 0.021 | 0.073 | 1.021 | C1,C2 |
| | *M. peruvianus* | 3 | Ocean | suction | OS | finned | 3.32 | +0.19 | 0.017 | 0.028 | 0.098 | 1.63 | 0.010 | 0.026 | 0.060 | 0.386 | C1,C2 |
| Lipotidae | *L. vexillifer* | 4 | River | bite | RB | finned | 2.20 | +0.23 | 0.018 | 0.022 | — | 0.64 | 0.028 | 0.052 | — | 0.538 | C2 |
| Iniidae | *I. geoffrensis* | 68 | River | bite | RB | finned | 2.03 | +0.18 | 0.001 | 0.036 | 0.035 | 0.55 | 0.002 | 0.101 | 0.064 | 0.022 | J,C2 |
| Pontoporiidae | *P. blainvillei* | 79 | River | bite | RB | finned | 1.32 | +0.24 | 0.007 | 0.009 | — | 0.23 | 0.031 | 0.060 | — | 0.509 | J,O,C1,C2 |

| | | | | | | | | | | | | | | | | | |
|---|---|---|---|---|---|---|---|---|---|---|---|---|---|---|---|---|---|
| Monodontidae | *D. leucas* | 2 | River | suction | RS | finless | 4.41 | ∅ | ∅ | 0.081 | 0.20 | 2.77 | ∅ | 0.045 | 0.074 | 0 | J,O,C1,C2 |
| | *M. monoceros* | 133 | River | suction | RS | finless | 4.33 | ∅ | ∅ | 0.043 | — | 2.68 | ∅ | 0.024 | — | 0 | J,C1,C2 |
| Phocoenidae | *Neophaena* sp. | 263 | River | bite | RB | finless | 1.32 | ∅ | ∅ | 0.013 | 0.020 | 0.30 | ∅ | 0.068 | 0.067 | 0 | O,C2 |
| | *P. phocoena* | 101 | Ocean | bite | OB | finned | 1.49 | +0.15 | 0.012 | 0.009 | 0.021 | 0.38 | 0.032 | 0.036 | 0.056 | 0.891 | O,C2 |
| | *P. dalli* | 212 | Ocean | bite | OB | finned | 1.90 | +0.07 | 0.033 | 0.008 | 0.070 | 0.62 | 0.054 | 0.021 | 0.113 | 2.596 | O,C2 |
| | *P. sinus* | 19 | Ocean | bite | OB | finned | 1.39 | +0.16 | 0.016 | 0.014 | 0.023 | 0.33 | 0.050 | 0.065 | 0.070 | 0.767 | C2 |
| | *P. dioptrica* | 16 | Ocean | bite | OB | finned | 1.99 | +0.18 | 0.037 | 0.009 | 0.031 | 0.67 | 0.055 | 0.020 | 0.046 | 2.747 | C2 |
| | *P. spinipinnis* | 11 | Ocean | bite | OB | finned | 1.66 | +0.31 | 0.016 | 0.015 | 0.024 | 0.47 | 0.035 | 0.049 | 0.051 | 0.714 | C2 |
| Delphinidae | *O. orca* | 6 | Ocean | bite | OB | finned | 5.77 | +0.05 | 0.36 | 0.35 | 0.48 | 3.63 | 0.099 | 0.146 | 0.132 | 0.676 | J,O,C1,C2 |
| | *O. brevirostris* | 23 | River | bite | RB | finned | 2.18 | +0.18 | 0.006 | 0.031 | 0.051 | 0.52 | 0.011 | 0.091 | 0.098 | 0.121 | C2 |
| | *O. heinsohni* | 20 | River | bite | RB | finned | 2.05 | +0.17 | 0.008 | 0.023 | — | 0.46 | 0.017 | 0.078 | — | 0.211 | C2 |
| | *G. griseus* | 17 | Ocean | suction | OS | finned | 2.50 | +0.16 | 0.069 | 0.053 | 0.068 | 0.68 | 0.102 | 0.118 | 0.100 | 0.857 | J,O,C1,C2 |
| | *P. crassidens* | 76 | Ocean | bite | OB | finned | 4.31 | +0.11 | 0.077 | 0.061 | 0.12 | 2.03 | 0.038 | 0.046 | 0.058 | 0.818 | O,C1,C2 |
| | *F. attenuata* | 12 | Ocean | bite | OB | finned | 2.26 | +0.15 | 0.042 | 0.036 | 0.041 | 0.56 | 0.076 | 0.099 | 0.074 | 0.766 | O,C1,C2 |
| | *P. electra* | 5 | Ocean | bite | OB | finned | 2.54 | +0.15 | 0.048 | 0.031 | 0.052 | 0.71 | 0.069 | 0.068 | 0.074 | 1.013 | C2 |
| | *G. macrorhynchus* | 102 | Ocean | bite | OB | finned | 3.88 | +0.03 | 0.11 | 0.066 | 0.14 | 1.64 | 0.068 | 0.062 | 0.085 | 1.098 | J,C2 |
| | *G. melas* | 99 | Ocean | bite | OB | finned | 4.32 | +0.11 | 0.095 | 0.12 | — | 2.03 | 0.047 | 0.089 | — | 0.527 | J,C1,C2 |
| | *L. albirostris* | 72 | Ocean | bite | OB | finned | 2.55 | +0.20 | 0.075 | 0.037 | — | 0.71 | 0.105 | 0.081 | — | 1.306 | C1,C2 |
| | *L. borealis* | 12 | Ocean | bite | OB | finless | 2.05 | ∅ | ∅ | 0.012 | 0.022 | 0.46 | ∅ | 0.041 | 0.047 | 0 | O,C1,C2 |
| | *L. peronii* | 1 | Ocean | bite | OB | finless | 2.28 | ∅ | ∅ | 0.014 | 0.029 | 0.57 | ∅ | 0.039 | 0.050 | 0 | C2 |
| | *C. commersonii* | 58 | Ocean | bite | OB | finned | 1.32 | +0.22 | 0.012 | 0.008 | 0.018 | 0.19 | 0.061 | 0.066 | 0.093 | 0.924 | J,C1,C2 |
| | *C. eutropia* | 13 | Ocean | bite | OB | finned | 1.51 | +0.23 | 0.016 | 0.010 | 0.020 | 0.25 | 0.064 | 0.061 | 0.081 | 1.042 | C1,C2 |
| | *L. obscurus* | 224 | Ocean | bite | OB | finned | 1.88 | +0.22 | 0.029 | 0.020 | 0.030 | 0.39 | 0.074 | 0.078 | 0.077 | 0.947 | C1,C2 |
| | *L. obliquidens* | 43 | Ocean | bite | OB | finned | 1.81 | +0.21 | 0.028 | 0.020 | 0.032 | 0.36 | 0.078 | 0.085 | 0.091 | 0.920 | O,C2 |
| | *C. heavisidii* | 13 | Ocean | bite | OB | finned | 1.61 | +0.17 | 0.021 | 0.010 | 0.025 | 0.28 | 0.074 | 0.054 | 0.089 | 1.364 | C1,C2 |
| | *L. australis* | 24 | Ocean | bite | OB | finned | 1.93 | +0.21 | 0.023 | 0.019 | 0.043 | 0.41 | 0.057 | 0.072 | 0.105 | 0.787 | C1,C2 |
| | *L. cruciger* | 2 | Ocean | bite | OB | finned | 1.77 | +0.19 | 0.026 | 0.017 | 0.034 | 0.34 | 0.076 | 0.078 | 0.100 | 0.970 | C1,C2 |
| | *S. bredanensis* | 3 | Ocean | bite | OB | finned | 2.45 | +0.14 | 0.038 | 0.031 | 0.055 | 0.66 | 0.058 | 0.072 | 0.083 | 0.803 | O,C1,C2 |
| | *S. fluviatilis* | 14 | Ocean | bite | OB | finned | 1.45 | +0.41 | 0.012 | 0.013 | 0.024 | 0.23 | 0.053 | 0.086 | 0.106 | 0.613 | C1 |
| | *S. plumbea* | 45 | Ocean | bite | OB | finned | 2.22 | +0.19 | 0.054 | 0.022 | — | 0.54 | 0.101 | 0.062 | — | 1.635 | C1,C2 |
| | *S. teuszii* | 6 | Ocean | bite | OB | finned | 2.23 | +0.13 | 0.063 | 0.021 | — | 0.54 | 0.116 | 0.060 | — | 1.932 | C2 |
| | *L. hosei* | 14 | Ocean | bite | OB | finned | 2.33 | +0.07 | 0.021 | 0.012 | 0.035 | 0.59 | 0.036 | 0.032 | 0.06 | 1.129 | O,C2 |
| | *S. longirostris* | 59 | Ocean | bite | OB | finned | 1.71 | +0.17 | 0.023 | 0.009 | 0.020 | 0.32 | 0.073 | 0.044 | 0.064 | 1.637 | O,C1,C2 |
| | *S. attenuata* | 125 | Ocean | bite | OB | finned | 1.93 | +0.18 | 0.024 | 0.009 | 0.025 | 0.41 | 0.059 | 0.032 | 0.060 | 1.835 | O,C1,C2 |
| | *T. aduncus* | 3 | Ocean | bite | OB | finned | 2.33 | +0.18 | 0.050 | 0.030 | 0.052 | 0.59 | 0.084 | 0.079 | 0.089 | 1.065 | C2 |
| | *T. truncatus* | 82 | Ocean | bite | OB | finned | 2.67 | +0.17 | 0.045 | 0.035 | 0.067 | 0.78 | 0.058 | 0.069 | 0.086 | 0.841 | J,O,C1,C2 |
| | *S. chinensis* | 10 | Ocean | bite | OB | finned | 2.18 | +0.20 | 0.030 | 0.021 | 0.054 | 0.52 | 0.058 | 0.062 | 0.105 | 0.931 | C2 |
| | *S. clymene* | 10 | Ocean | bite | OB | finned | 1.86 | +0.13 | 0.022 | 0.011 | 0.026 | 0.38 | 0.058 | 0.046 | 0.070 | 1.278 | C1,C2 |
| | *S. coeruleoalba* | 50 | Ocean | bite | OB | finned | 2.07 | +0.17 | 0.024 | 0.013 | 0.031 | 0.47 | 0.052 | 0.044 | 0.066 | 1.187 | O,C1,C2 |
| | *S. frontalis* | 79 | Ocean | bite | OB | finned | 1.97 | +0.21 | — | 0.013 | 0.034 | 0.42 | — | 0.047 | 0.08 | — | C1 |
| | *D. delphis* | 103 | Ocean | bite | OB | finned | 2.06 | +0.19 | 0.032 | 0.017 | 0.032 | 0.46 | 0.068 | 0.057 | 0.068 | 1.199 | J,O,C1,C2 |

**Note:** $a^*_{T,\mathrm{lat}}$ was estimated using the following family-level fineness ratios [FR] based on Fish and Rohr (1999): Delphinidae, 7.2; Phocoenidae, 4.6; Monodontidae, 5.5; Platanistidae, 5.9; Ziphiidae, 5.3; Physeteridae, 5.9; Balaenopteridae, 6.3; and Balaenidae, 5.3. Ecotype abbreviations were as follows: OF, oceanic filter feeder; OS, oceanic suction feeder; OB, oceanic bite feeder; RS, riverine suction feeder; and RB, riverine bite feeder. Ref. (FB) were as follows: (J) Johnston, C., and Berta, A. (2011) Comparative anatomy and evolutionary history of suction feeding in cetaceans. *Marine Mammal Science* 27, 493–513; (O) Okamura, T., and Fujiwara, S. (2020) The range of atlanto-occipital joint motion in cetaceans reflects their feeding behavior. *Journal of Anatomy* 236, 434–447; (C1) Churchill, M., and Baltz, C. (2021) Evolution of orbit size in toothed whales (Artiodactyla: Odontoceti). *Journal of Anatomy* 239, 1419–1437; and (C2) Coombs, E. J., Felice, R. N., Clavel, J., Park, T., Bennion, R. F., Churchill, M., Geisler, J. H., Beatty, B., and Goswami, A. (2022) The tempo of cetacean cranial evolution. *Current Biology* 32, 2233–2247.

**Table S3** Steel–Dwass comparison results for $DFR^*$, $A_D^*$, $A_F^*$, and $A_C^*$ among cetacean ecotypes.

| pairs | $DFR^*$ | | $A_D^*$ | | $A_F^*$ | | $A_C^*$ | |
|---|---|---|---|---|---|---|---|---|
| | t | p | t | p | t | p | t | p |
| OF : OS | −5.62 | 4.09E-04* | −4.90 | 2.97E-03* | −4.15 | 1.75E-02* | −0.18 | 1.00 |
| OF : OB | −6.70 | 1.29E-05* | −6.70 | 1.29E-05* | −2.05 | 4.68E-01 | −1.34 | 0.78 |
| OF : RB | −0.84 | 9.32E-01 | −2.77 | 2.05E-01 | −2.63 | 2.44E-01 | −0.20 | 1.00 |
| OS : OB | −5.48 | 6.13E-04* | −6.71 | 1.23E-05* | −7.42 | 9.32E-07* | −2.14 | 0.43 |
| OS : RB | −3.88 | 3.07E-02* | −0.78 | 9.46E-01 | −5.05 | 2.04E-03* | −0.85 | 0.93 |
| OB : RB | −5.14 | 1.56E-03* | −5.19 | 1.37E-03* | −2.70 | 2.25E-01 | −0.30 | 0.99 |

**Note:** OF, oceanic filter feeder; OS, oceanic suction feeder; OB, oceanic bite feeder; RB, river bite feeder. $t$, test statistic; $p$, adjusted $P$ value. Asterisks indicate significant differences ($P < 0.05$).

**Table S4** AIC values for discrete-trait evolutionary models fitted to dorsal fin presence (finned) and ecotype on the molecular phylogenies of McGowen et al. (2020) and McGowen et al. (2009).

| | M20 | | M09 | |
|---|---|---|---|---|
| Model | finned | Ecotype | finned | Ecotype |
| ER | 41.26 | 72.98 | 40.81 | 78.16 |
| SYM | 41.26 | 80.16 | 40.81 | 85.65 |
| ARD | 42.48 | 96.19 | 42.54 | 99.84 |

**Note:** ER, equal-rates model; SYM, symmetric-rates model; ARD, all-rates-different model; M20, McGowen et al. (2020) *Syst. Biol.* 69, 479–501; M09, McGowen et al. (2009) *Mol. Phylogenet. Evol.* 53, 891–906.
Lowest AIC values indicate best model fit.

**Table S5** AIC values for continuous-trait evolutionary models fitted to $A_D^*$, $A_F^*$, and $A_C^*$ on the phylogenies of M09 and M20.

| | M20 | | | M09 | | |
|---|---|---|---|---|---|---|
| Model | $A_D^*$ | $A_F^*$ | $A_C^*$ | $A_D^*$ | $A_F^*$ | $A_C^*$ |
| OU | −294.95 | −280.46 | −273.10 | −344.58 | −315.84 | −313.20 |
| BM | −290.09 | −277.31 | −270.32 | −337.15 | −313.78 | −304.06 |
| EB | −288.09 | −275.31 | −268.32 | −335.15 | −311.78 | −302.06 |

**Note:** OU, Ornstein–Uhlenbeck model; BM, Brownian motion model; EB, early-burst model.

**Table S6** AIC values for OUwie models fitted to $A_D^*$, $A_F^*$, and $A_C^*$ on the phylogenies of M09 and M20.

| | M20 | | | M09 | | |
|---|---|---|---|---|---|---|
| Model | $A_D^*$ | $A_F^*$ | $A_C^*$ | $A_D^*$ | $A_F^*$ | $A_C^*$ |
| OUM | −348.87 | (−280.46) | (−273.10) | −364.21 | −341.77 | (−313.20) |
| OU1 | −299.52 | (−280.46) | (−273.10) | −344.58 | −315.84 | (−313.20) |
| BM1 | −290.09 | (−277.31) | (−270.32) | −337.15 | −313.78 | (−304.06) |

**Note:** OUM, multiple-optimum OU model; OU1, single-optimum OU model; BM1, single-rate BM model.
Values in parentheses indicate traits for which no selective regime shift was detected within the phylogeny.

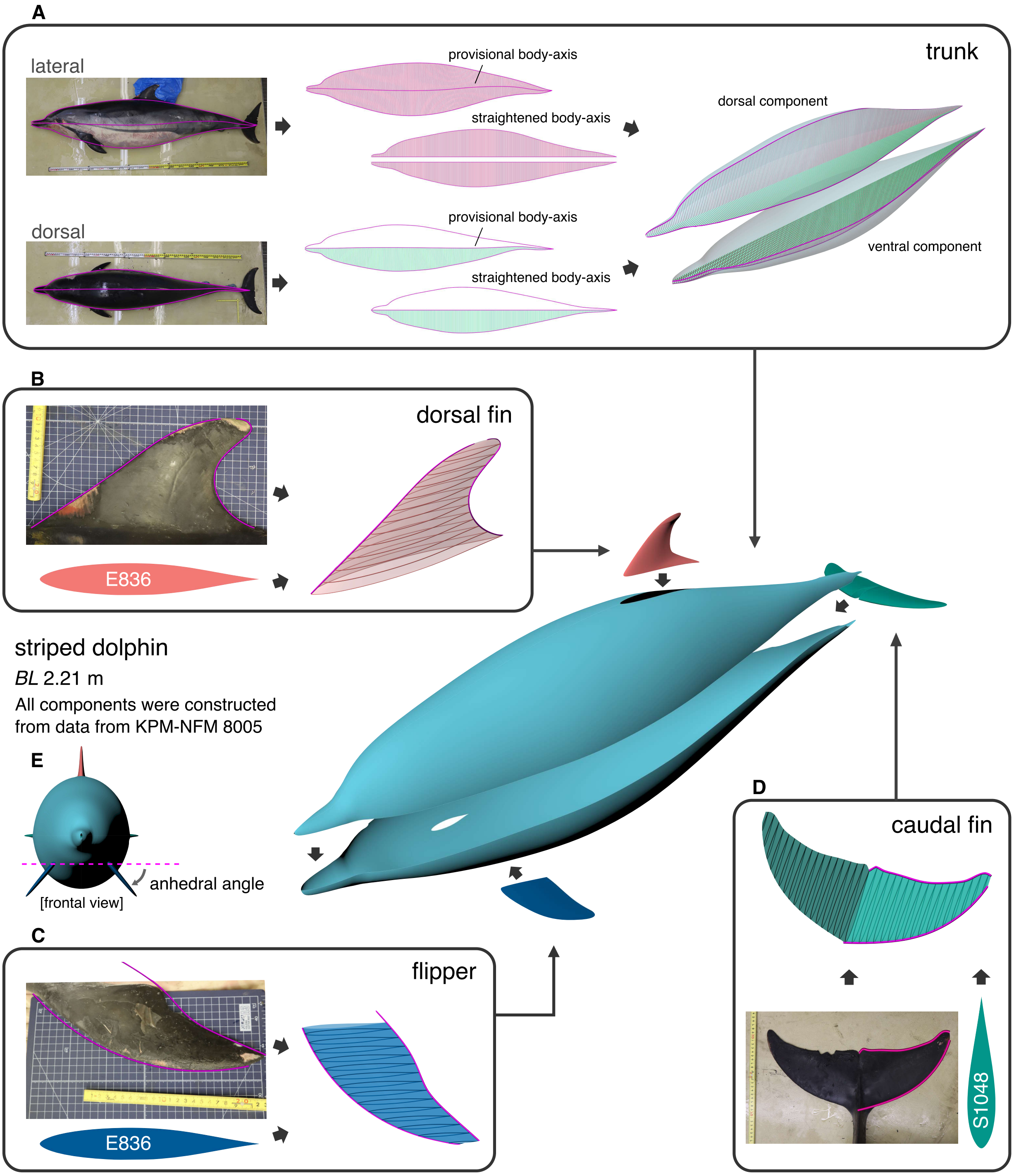


**Figure SM1 3D reconstruction workflow of the cetacean CFD model.**
The cetacean CFD model comprised four major components, the trunk (A), dorsal fin (B), flipper (C), and caudal fin (D), which were generated separately and then assembled into a full-body model. For example, the striped dolphin model was constructed from photographs and external morphometric measurements of a single individual stranded in Kanagawa, Japan (*BL* [body length] = 2.21 m; KPM-NFM 8005). The trunk was reconstructed by first geometrically correcting the outlines to straighten the anteroposterior axis and then separately generating dorsal and ventral components from half-elliptical cross sections fitted to the corrected outlines. The dorsal fin, flipper, and caudal fin were modeled separately by tracing their leading and trailing edges from photographs and reference images with NURBS curves and applying hydrofoil profiles approximating cetacean fins. The E836 profile was used for the flipper and dorsal fin, and the S1048 for the caudal fin (Pavlov et al., 2021). These components were then assembled with reference to the external morphometric measurements to generate the complete 3D model. All fins were fixed at an angle of attack of 0° under the 0° yaw condition. Only the flipper varied among simulation cases, with its anhedral angle changed by rotation about the shoulder joint while maintaining the minimum sweep configuration (E). Fin shapes were locally adjusted while preserving fin area to ensure seamless junctions between adjacent components.

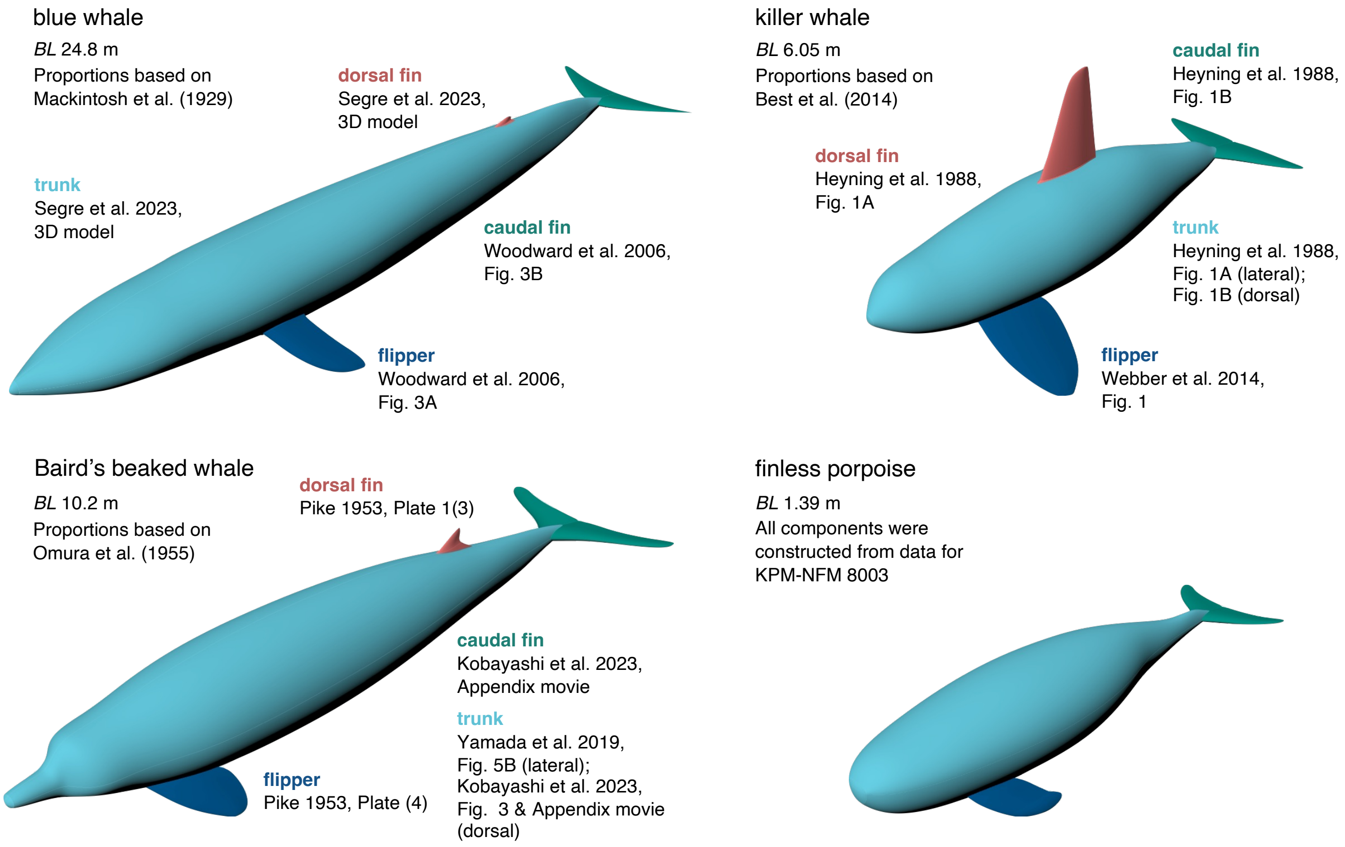


**Figure SM2 CFD models of the other four species and the source data used for their construction.** The blue whale, Baird's beaked whale, and killer whale models were constructed by generating body components from image data of multiple individuals reported in previous studies, scaling them according to body proportions derived from external morphometric measurements, and assembling them into a single full-body model. The finless porpoise model, like the striped dolphin model, was constructed from photographs and external morphometric measurements of an individual stranded in Kanagawa, Japan (KPM-NFM 8003). The references for the source data are listed below.

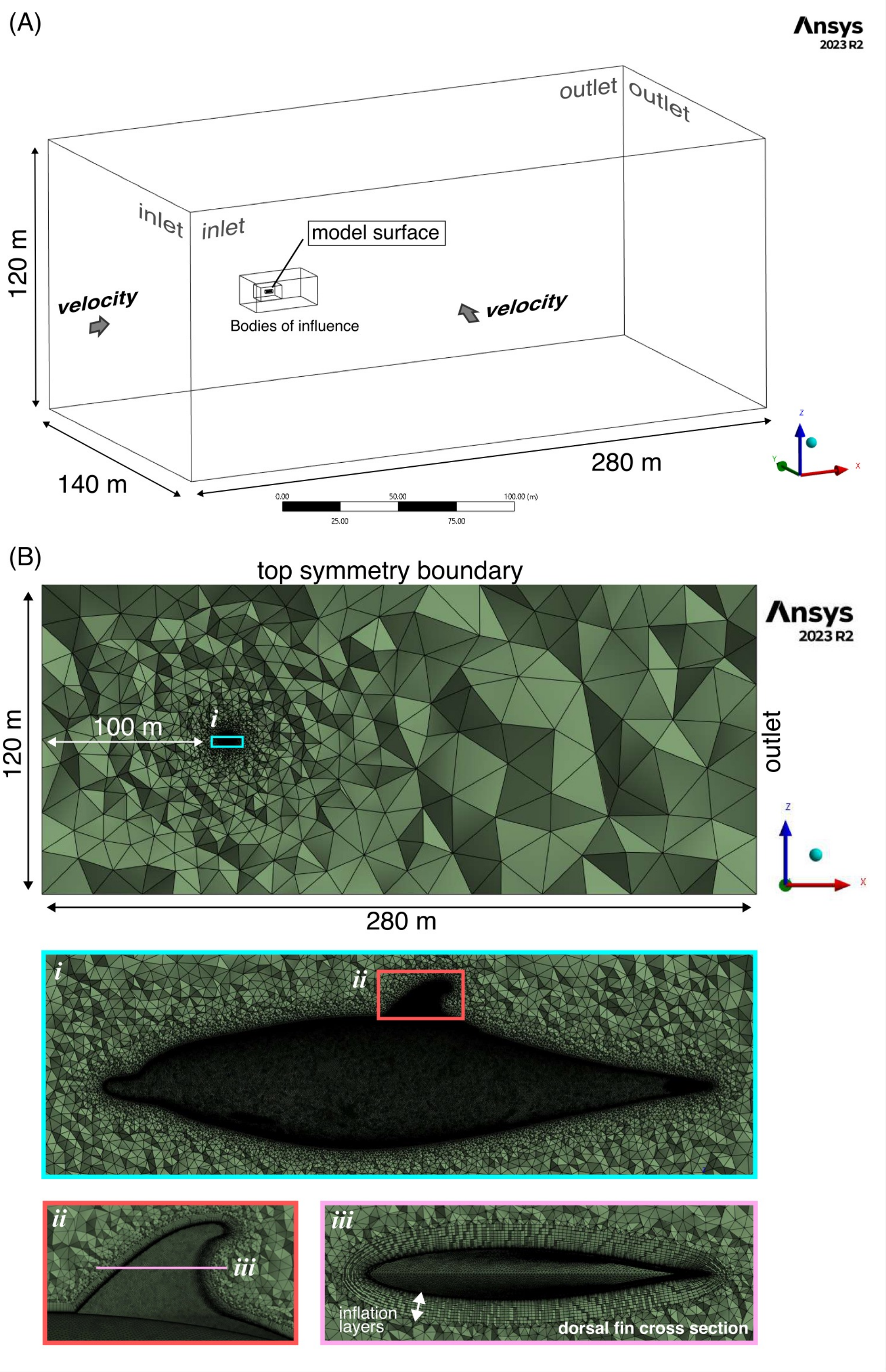


**Figure SM3 Computational domain and mesh configuration used for the CFD simulations.**
(A) Computational domain and boundary conditions used for the CFD simulations. The cetacean model was placed in a sufficiently large cuboidal fluid domain (280 m × 140 m × 120 m in the X-, Y-, and Z-directions, respectively) to minimize boundary effects. Boundary conditions consisted of a velocity inlet at the upstream face, a pressure outlet at the downstream face, symmetry conditions on the top and bottom surfaces, and a no-slip wall on the model surface. At the lateral boundaries, the upstream-side boundary in the Y-direction was defined as a velocity inlet and the opposite-side boundary as a pressure outlet. (B) Mesh configuration shown at four levels of magnification: the entire computational domain, the whole-body mesh of the striped dolphin model (*i*), an enlarged view of the dorsal fin region (*ii*), and a cross-sectional view of the dorsal fin (*iii*). The mesh consisted primarily of tetrahedral elements, with local refinement around the model achieved using nested Body of Influence regions. A boundary-layer mesh consisting of 20 prism layers was applied to the model surface, with the first-layer thickness adjusted to maintain $y^+ < 5$, and the inflation layers are visible in the dorsal fin cross section (*iii*).

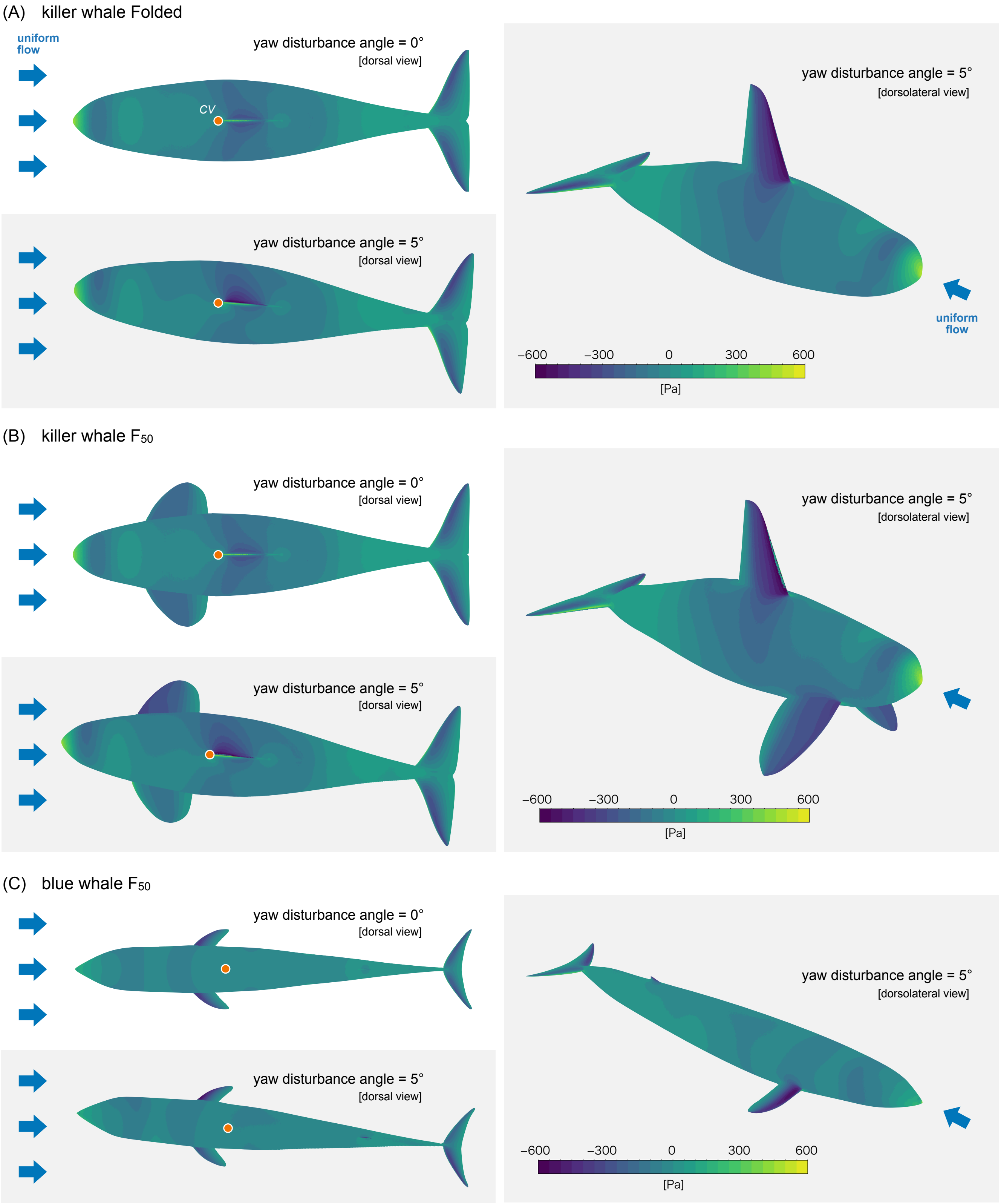


**Figure SM4 Pressure distributions under yaw disturbance.**
Pressure distributions in dorsal and dorsolateral views for (A) the killer whale model with folded flippers (killer whale folded), (B) the killer whale model with the flippers extended at a 50-degree anhedral angle (killer whale $F_{50}$), and (C) the blue whale model with the flippers extended at a 50-degree anhedral angle (blue whale $F_{50}$), at an inflow velocity of 1 m $s^{-1}$ under no yaw disturbance (0° yaw disturbance) and under a 5° yaw disturbance. The center of volume ($CV$) is overlaid in the dorsal views. Reynolds numbers were $5.3\times10^6$ for the killer whale models, $21.8\times10^6$ for the blue whale model. Under no yaw disturbance, the pressure distributions were bilaterally symmetrical. Under the 5° yaw disturbance, marked left-right asymmetry appeared, particularly around the dorsal fin and flippers of the killer whale and the flippers of the blue whale, indicating lift generation on these appendages. However, the resulting yawing moments remained very small because these structures were located close to $CV$.

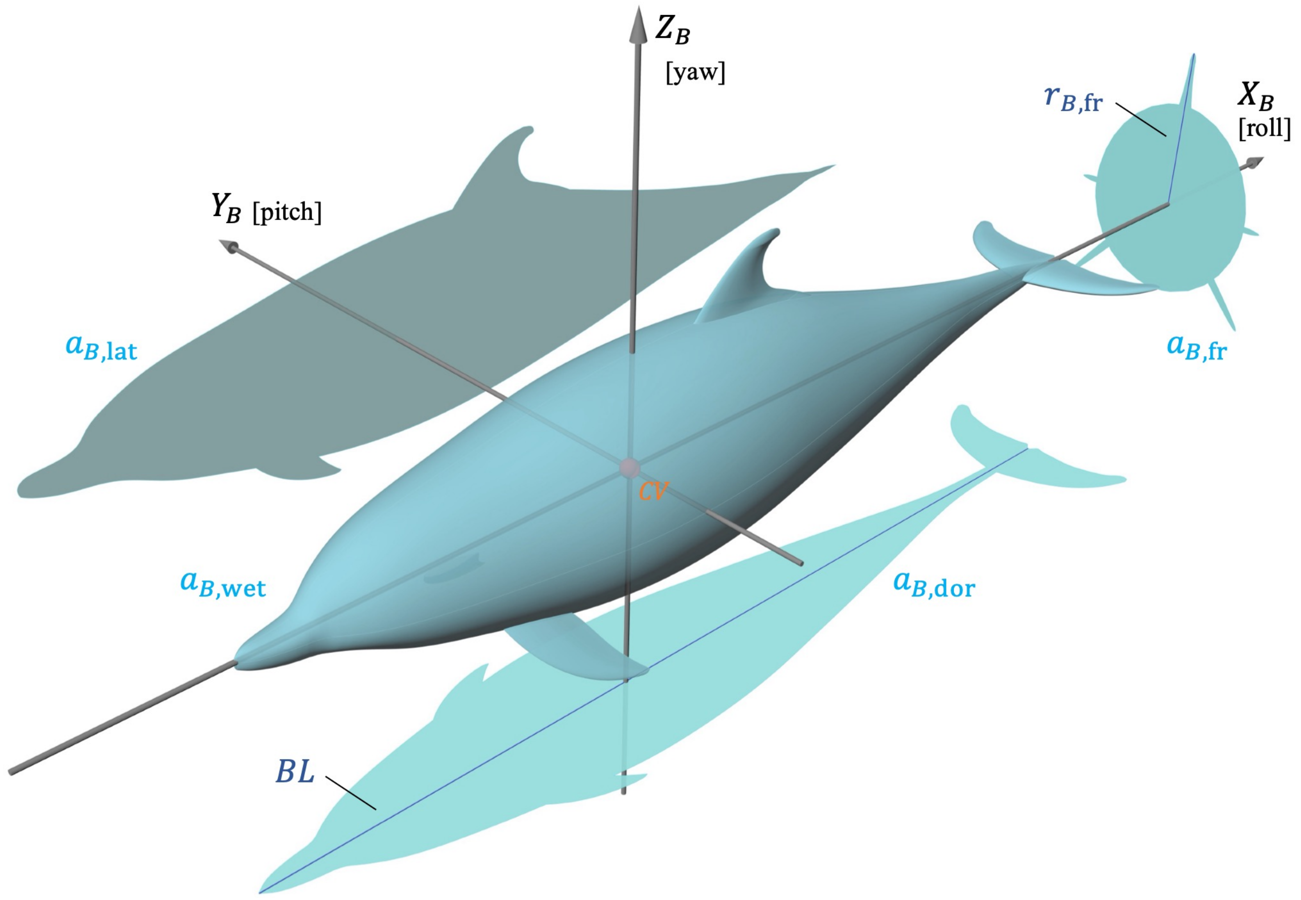


**Figure SM5 Body axes of the CFD model and variables used to normalize CFD simulation results.**
The body axes ($X_B$, $Y_B$, $Z_B$) were defined with the center of volume ($CV$) as the center of rotation. $a_{B,\mathrm{wet}}$ is the wetted surface area of the whole body; $a_{B,\mathrm{dor}}$ is the dorsal projected area of the whole body; BL is body length; $a_{B,\mathrm{lat}}$ is the lateral projected area of the whole body; $a_{B,\mathrm{fr}}$ is the frontal projected area of the whole body; and $r_{B,\mathrm{fr}}$ is the maximum distance from the body axis in the frontal projection of the whole body. In the dolphin $F_{50}$ model shown here, $r_{B,\mathrm{fr}}$ corresponds to the distance from the body axis to the tip of the dorsal fin, although the farthest point varied among models and could occur at the flipper tip, trunk, or other body parts. These distances and areas were measured using Rhinoceros.

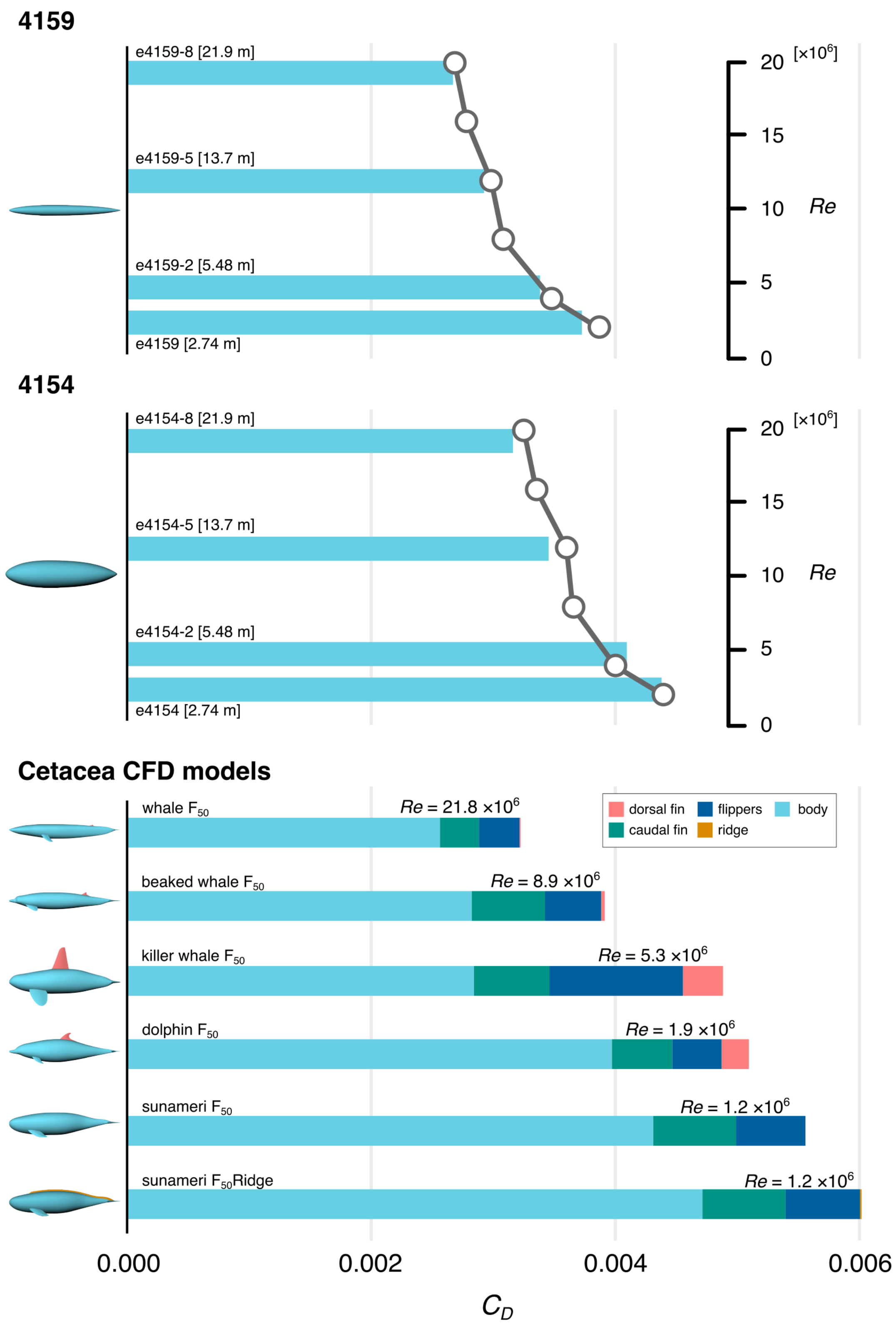


**Figure SM6 Drag-size validation using prolate spheroid models and cetacean CFD models.** Drag coefficients ($C_D$) obtained by CFD for the prolate spheroid models 4154 and 4159 are shown as bar plots at Reynolds numbers ($Re$) corresponding to geometrically scaled model lengths of 2.74, 5.48, 13.7, and 21.9 m ($Re$ = $2.4\times10^6$, $4.8\times10^6$, $12.0\times10^6$, and $19.2\times10^6$, respectively). Experimental values reported by Gertler (1950) are overlaid as points for comparison. The CFD-derived $C_D$ of the prolate spheroids agreed closely with the experimental values, with deviations within 5%. Drag coefficients of the cetacean CFD models are also shown at their corresponding Reynolds numbers, with the contributions of each body component displayed separately.

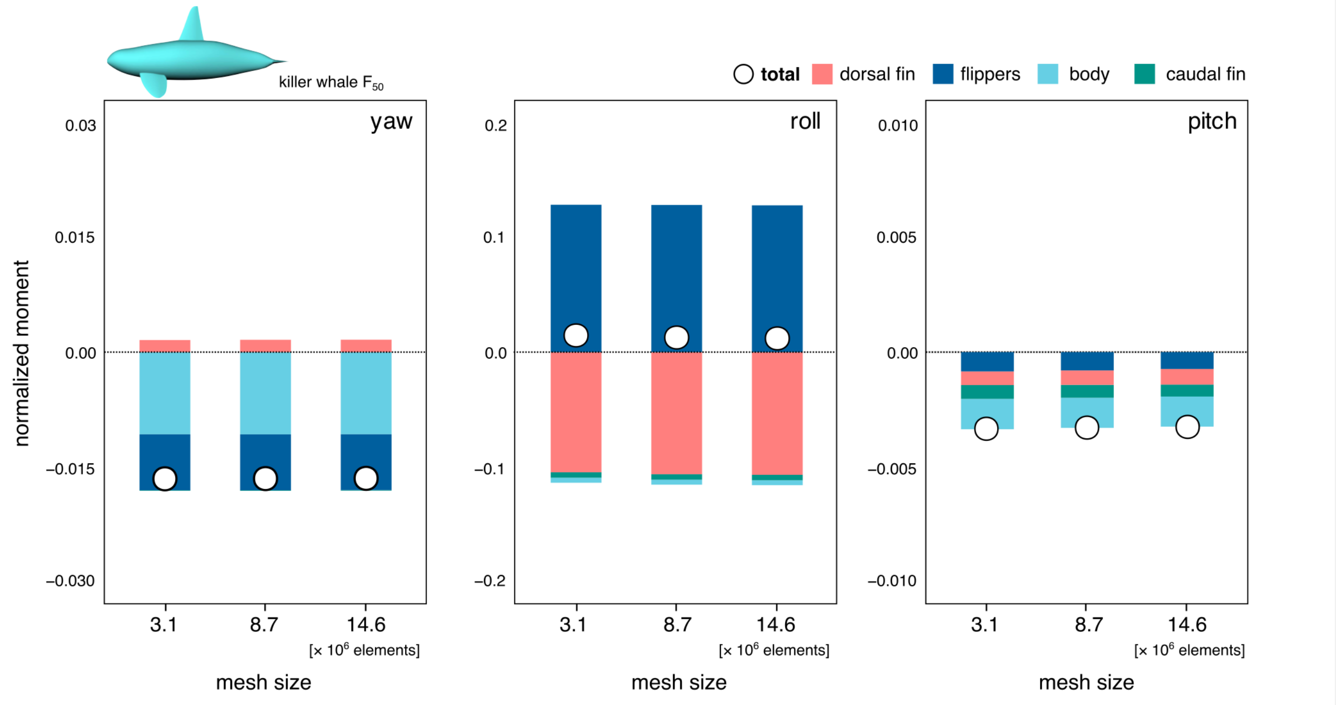


**Figure SM7 Mesh-size validation in CFD simulations.**
Normalized yawing, rolling, and pitching moments of each body component, together with the normalized total moments, are shown for the killer whale model at flow speed of 1 m s$^{-1}$ under a 5° yaw disturbance for computational domains generated using meshes of 3.1, 8.7, and 14.6 million elements. Differences among mesh sizes were minimal, indicating that the results were robust to mesh resolution.

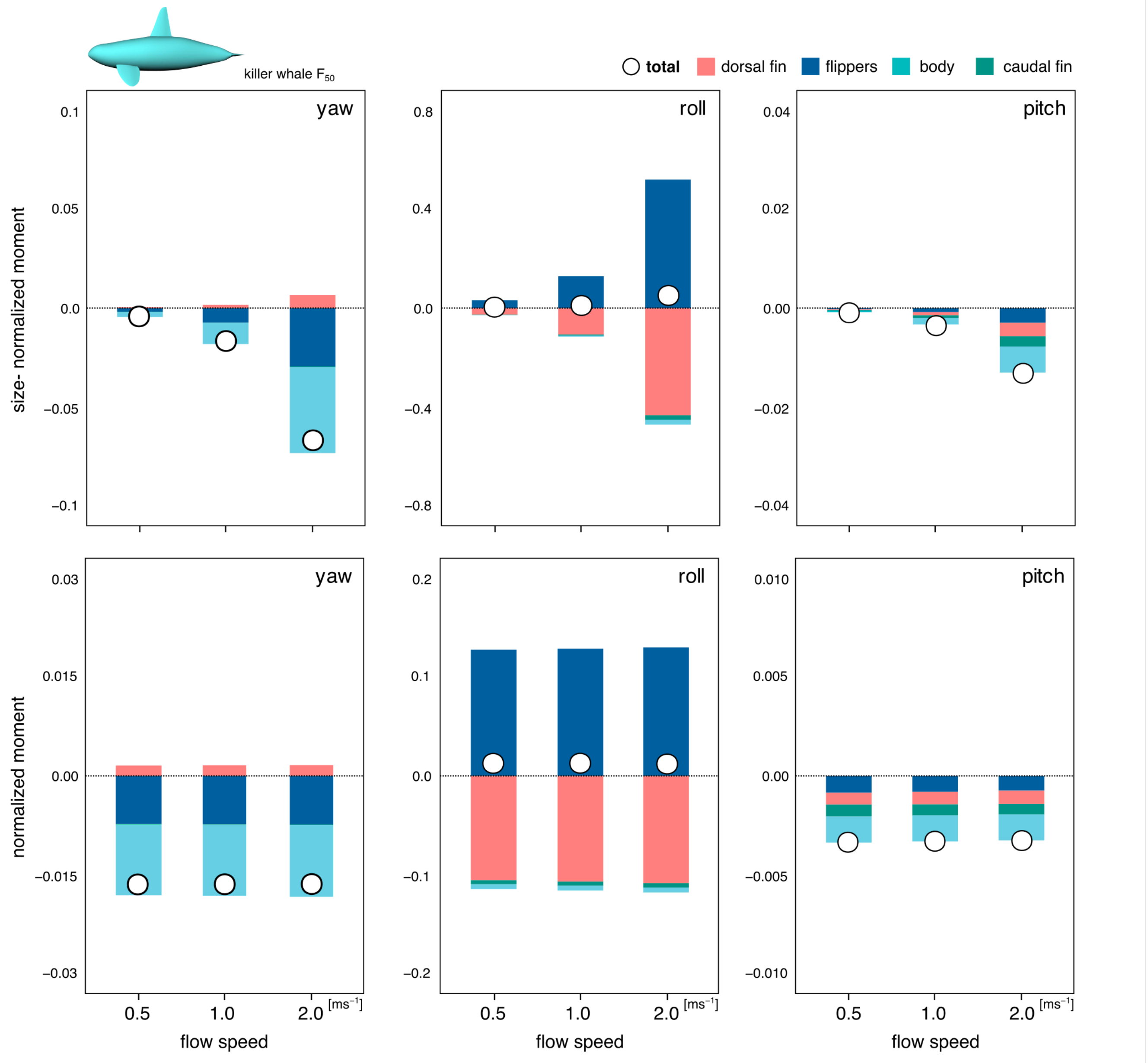


**Figure SM8 Effects of flow speed in CFD simulations.**
Normalized yawing, rolling, and pitching moments of each body component, together with the normalized total moments, are shown for the killer whale model under a 5° yaw disturbance at flow speeds of 0.5, 1.0, and 2.0 m s$^{-1}$. The upper panels show moments normalized by projected area and arm length only (size- normalized moment), whereas the lower panels show moments further normalized by dynamic pressure. Although moment magnitudes increased approximately in proportion to the square of flow speed, the fully normalized values varied little, indicating that moment behavior was largely independent of flow speed within the range of typical cetacean swimming speeds.

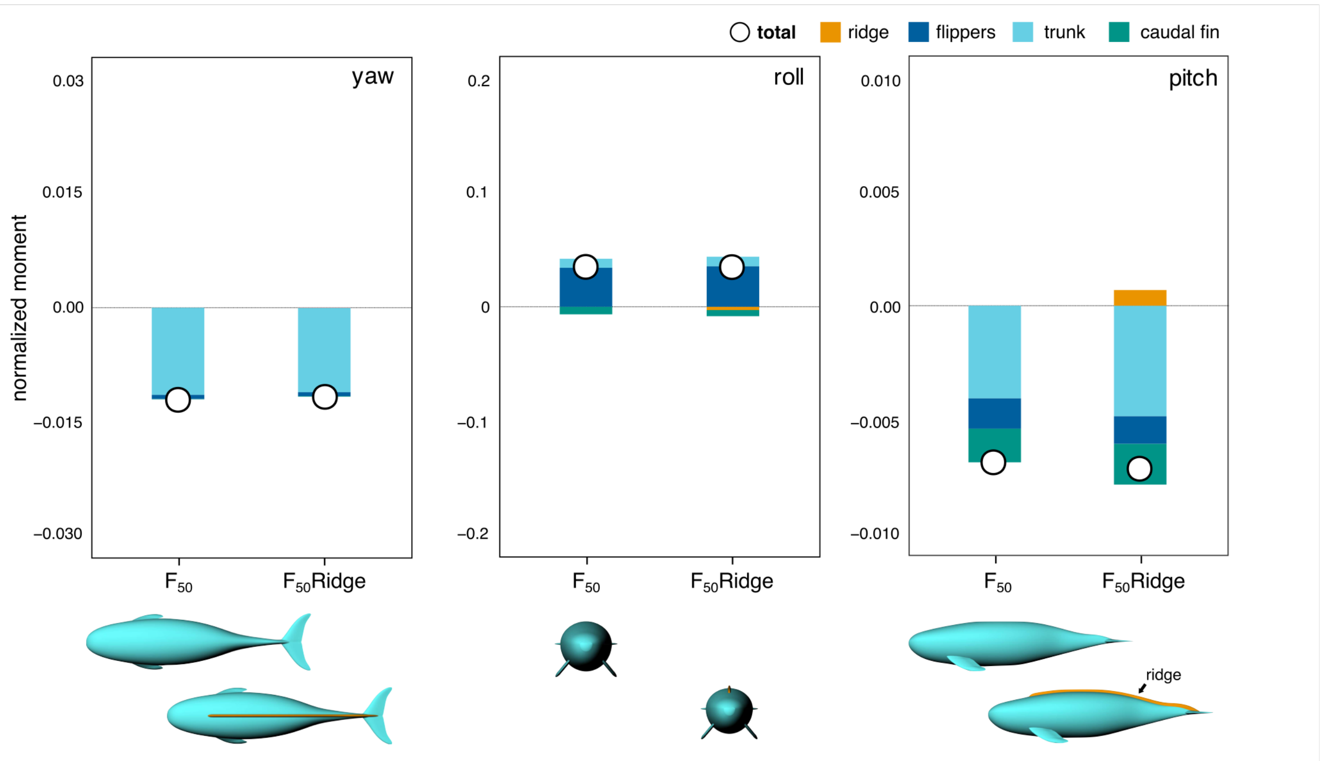


**Figure SM9 Effects of the dorsal ridge in the finless porpoise.**
Normalized yawing, rolling, and pitching of each body component, together with the whole-body normalized total moments, are shown for finless porpoise models with (sunameri $F_{50}$) and without the dorsal ridge (sunameri $F_{50}$ Ridge) at flow speed of 1 m s$^{-1}$ under a 5° yaw disturbance angle. Ridge effects were small and did not alter the overall trends observed in the main study.

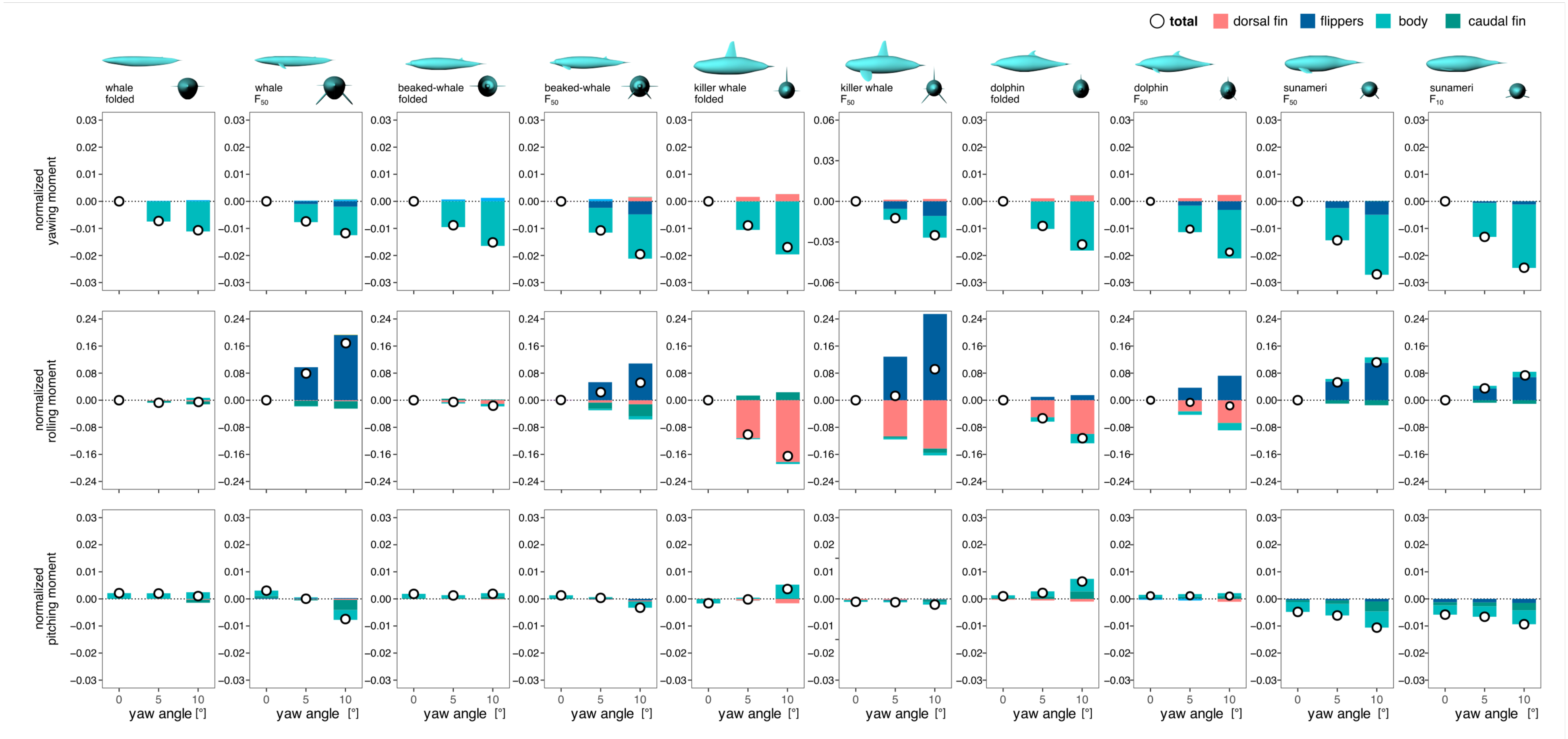


**Figure SR1 Moments of each body component across yaw disturbance angles in each CFD model.**
For each CFD model, normalized yawing, rolling, and pitching moments of each body component, together with the whole-body normalized total moments, are shown at flow speed of 1 m s$^{-1}$ under yaw disturbance angles of 0°, 5°, and 10°. The vertical axis shows moment magnitude, and the horizontal axis shows yaw disturbance angle.

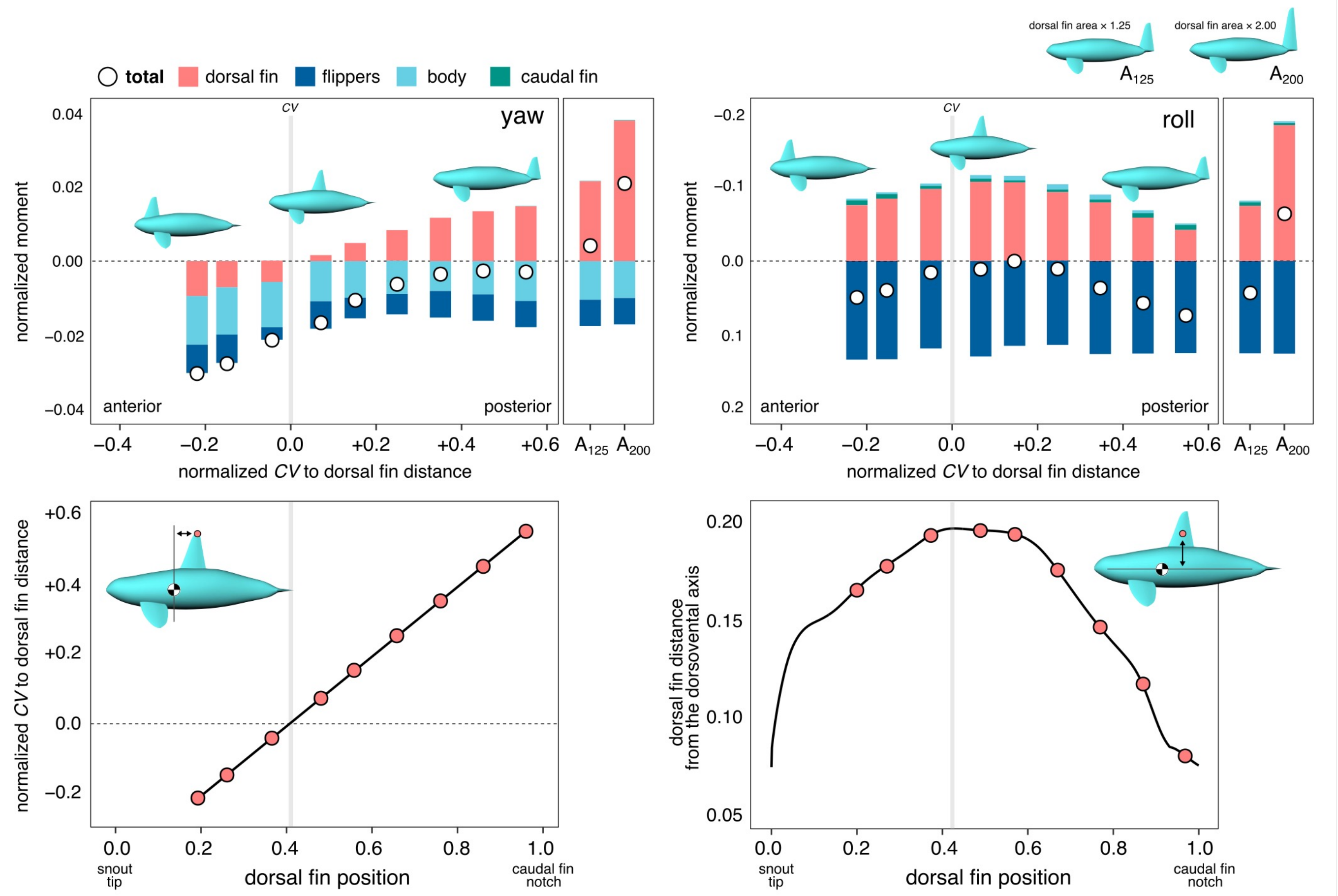


**Figure SR2 Effects of dorsal fin position and area in hypothetical killer whale models.**
Normalized yawing and rolling moments are shown for hypothetical killer whale models with altered dorsal fin position and area at flow speed of 1 m s⁻¹ under a 5° yaw disturbance. In the position-modified models, dorsal fin position was varied while dorsal fin area was held constant. The upper panels show the normalized moments of each body component together with the normalized total moments. The vertical axis shows normalized moment magnitude, and the horizontal axis shows the normalized distance from the center of rotation ($CV$) to the dorsal fin tip (defined $DP_C$) and model name ($A_{125}$ and $A_{200}$). $DP_C$ ranges from −0.21 to +0.54, with negative and positive values indicating dorsal fin positions anterior and posterior to the center of rotation, respectively. The highlighted model indicates the dorsal fin position of the real killer whale ($DP_C$ = +0.08). $A_{125}$ and $A_{200}$ denote models in which the dorsal fin was placed at the most posterior position ($DP_C$ = +0.54) and enlarged to 1.25 and 2.00 times the original area, respectively. The lower panels show the distances from the body axis passing through the center of rotation to the dorsal fin. For yawing and rolling, these correspond to the anteroposterior and dorsoventral distances, respectively.

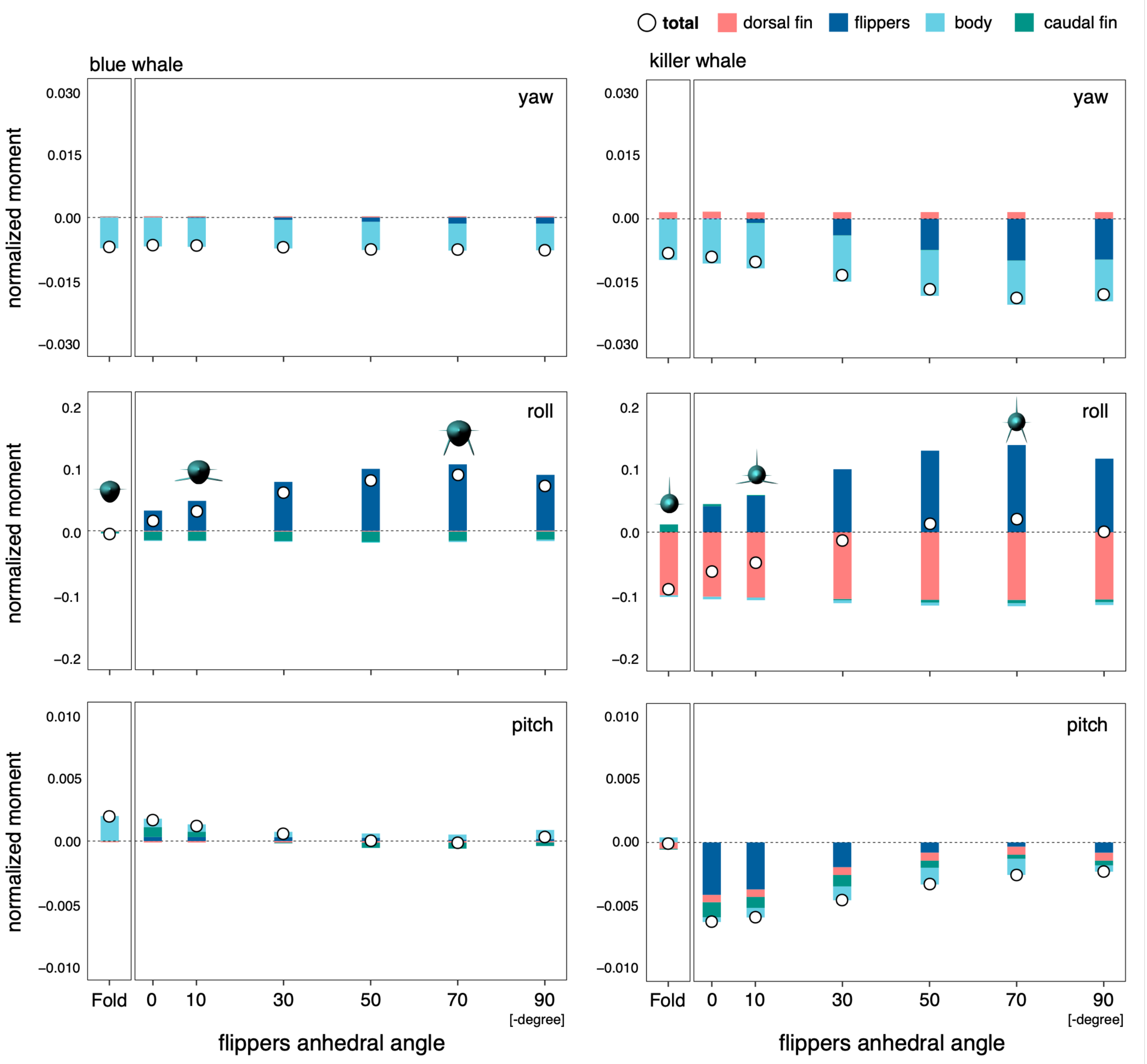


**Figure SR3 Effects of flipper posture in the blue whale and killer whale models.**
Normalized yawing, rolling, and pitching moments of each body component, together with the whole-body normalized total moments, are shown for the blue whale and killer whale models at flow speed of 1 m $s^{-1}$ under a 5° yaw disturbance, with the flippers either folded or extended at 0-, 10-, 30-, 50-, 70-, and 90-degree anhedral angles. For the extended-posture models, flipper area was held constant across anhedral angles. The vertical axis shows normalized moment magnitude, and the horizontal axis shows flipper posture.

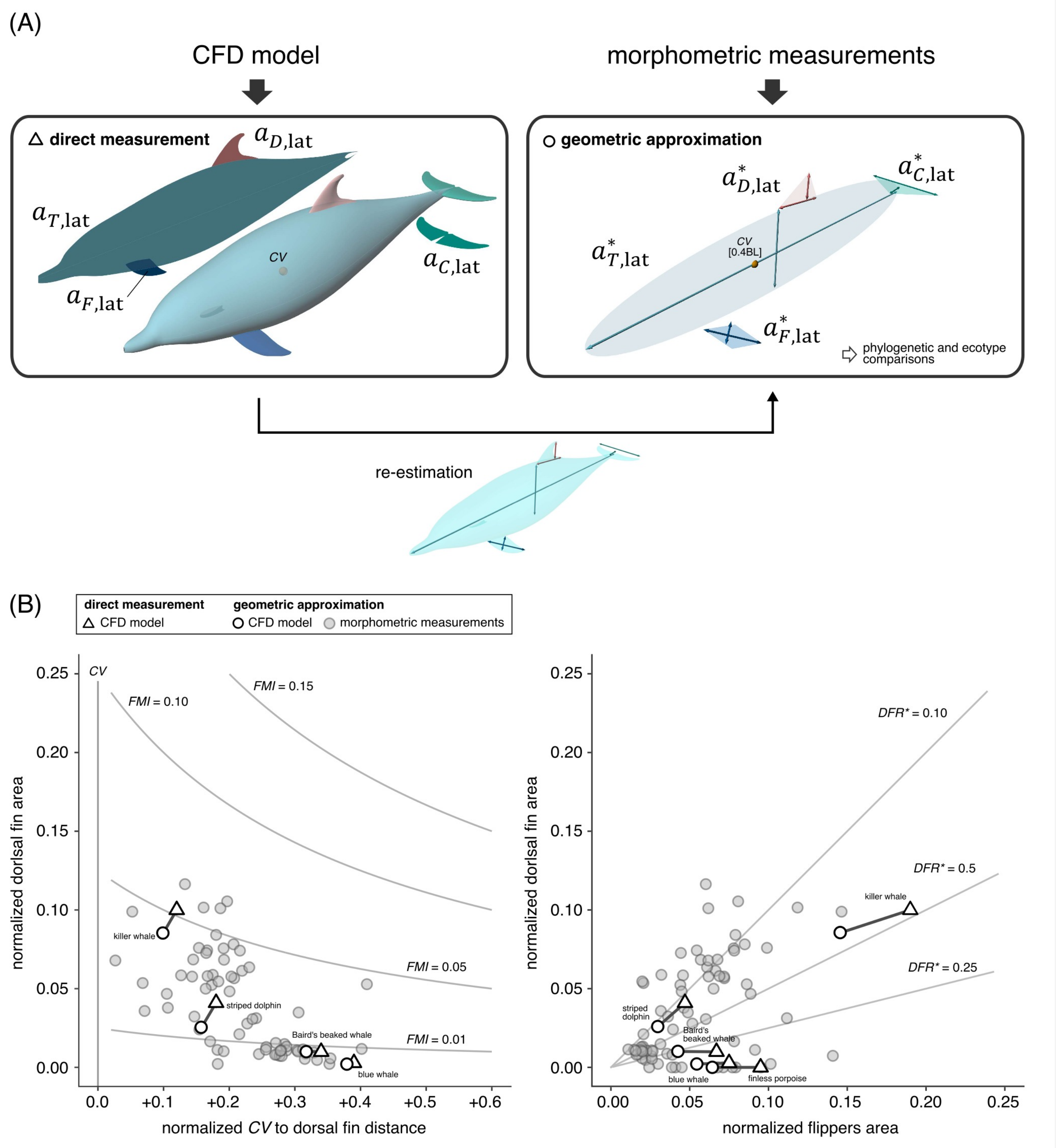


**Figure SC1 Projected area calculation by two methods and their distributions.**
(A) Projected areas of the fins and trunk were obtained by two methods: direct measurement from the CFD models, yielding $a_{D,\mathrm{lat}}$; $a_{F,\mathrm{lat}}$; $a_{C,\mathrm{lat}}$; $a_{T,\mathrm{lat}}$, and geometric approximation based on external morphometric measurements, yielding $a^*_{D,\mathrm{lat}}$; $a^*_{F,\mathrm{lat}}$; $a^*_{C,\mathrm{lat}}$; $a^*_{T,\mathrm{lat}}$, where asterisks indicate approximated values. In the latter, projected areas were estimated from two-point distance data by approximating the flipper as a rhombus, the dorsal and caudal fins as triangles, and the trunk as an ellipse. The center of volume ($CV$) was standardized at $0.4BL$ from the snout tip across species. To evaluate whether CFD-based results can be extended to interspecific analyses based on external morphometric data, both direct measurements and geometric re-estimates were obtained for the CFD models. (B) The distributions of values obtained for the five CFD models by direct measurement and geometric re-estimation are superimposed on those obtained for extant cetaceans by geometric approximation. These panels use the same x- and y-axes as Fig. 2A,B, respectively. See the main text for the calculation of $FMI$ and $DAR^*$. Although geometric estimates tended to be smaller than direct measurements, the discrepancies were small relative to the overall diversity of fin morphology across cetaceans and did not substantially alter the relative size relationship between the dorsal fin and the flippers.

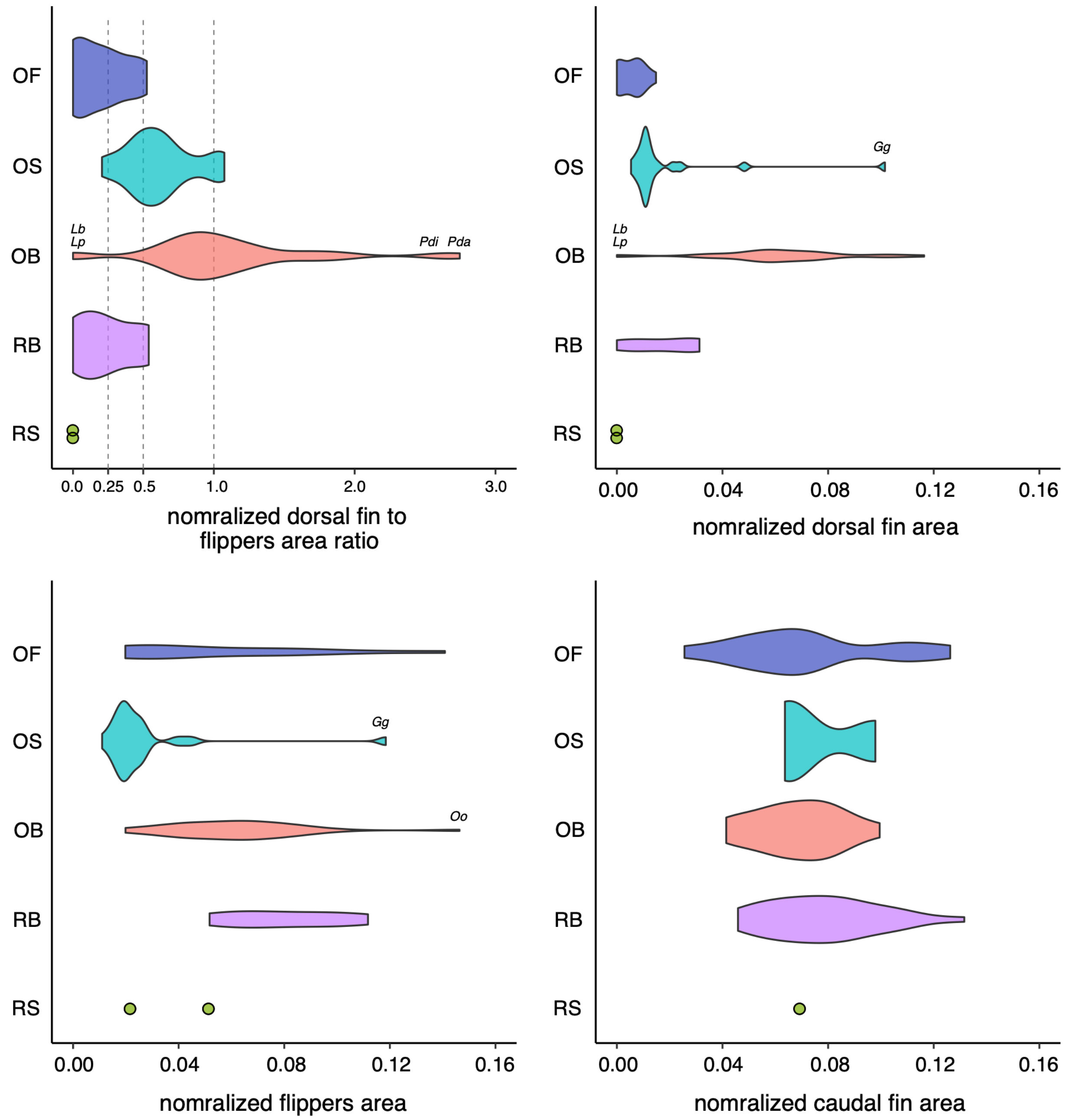


**Figure SC2 Violin plots of normalized fin area variables across ecotypes in extant cetaceans.**
Violin plots show the normalized dorsal fin to flippers area ratio ($DFR^*$), geometrically approximated normalized dorsal fin area ($A_D^*$), geometrically approximated normalized flippers area ($A_F^*$), and geometrically approximated normalized caudal fin area ($A_C^*$) across ecotypes in extant cetaceans: oceanic filter feeder (OF), oceanic suction feeder (OS), oceanic bite feeder (OB), riverine bite feeder (RB), and riverine suction feeder (RS). $A_D^*$ showed clearer ecotype-level differences than $A_F^*$ or $A_C^*$, producing corresponding differences in $DFR^*$. In particular, DFR* in OB was concentrated around 1.0, indicating that $A_D^*$ approached $A_F^*$ in this ecotype. Species names shown above the violins denote species with extreme values: *Lb*, *Lissodelphis borealis*; *Lp*, *Lissodelphis peronii*; *Gg*, *Grampus griseus*; *Oo*, *Orcinus orca*; *Pdi*, *Phocoena dioptrica*; and *Pda*, *Phocoenoides dalli*.

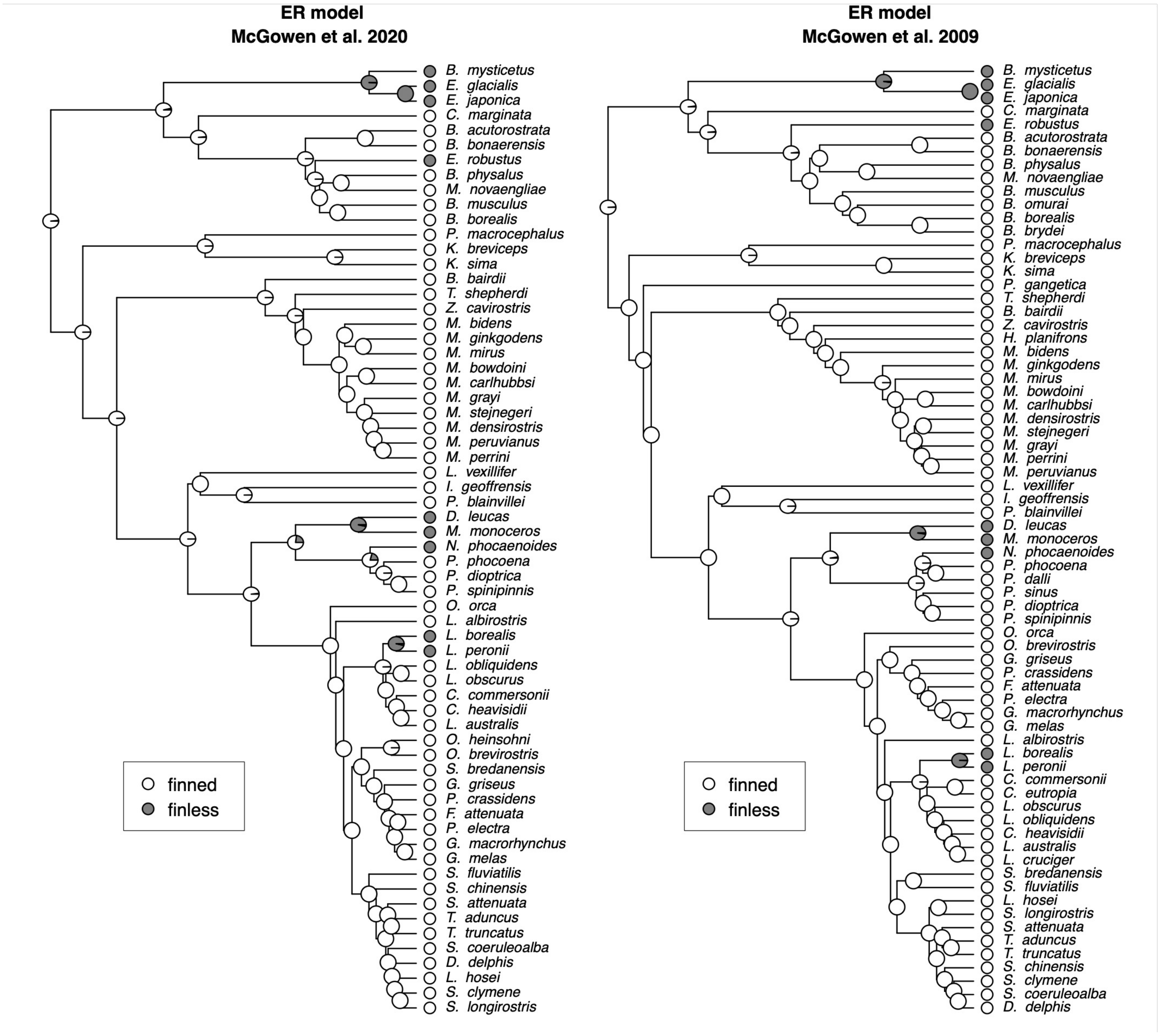


**Figure SC3 Ancestral state reconstruction of dorsal fin presence or absence.**
Ancestral state reconstruction of dorsal fin presence or absence (finned or finless) under the equal-rates (ER) model is shown for the molecular phylogenies of McGowen et al. (2020) and McGowen et al. (2009). In both phylogenies, the common ancestor of extant cetaceans had a very high probability of already possessing a dorsal fin, suggesting multiple independent losses of the dorsal fin in different lineages.

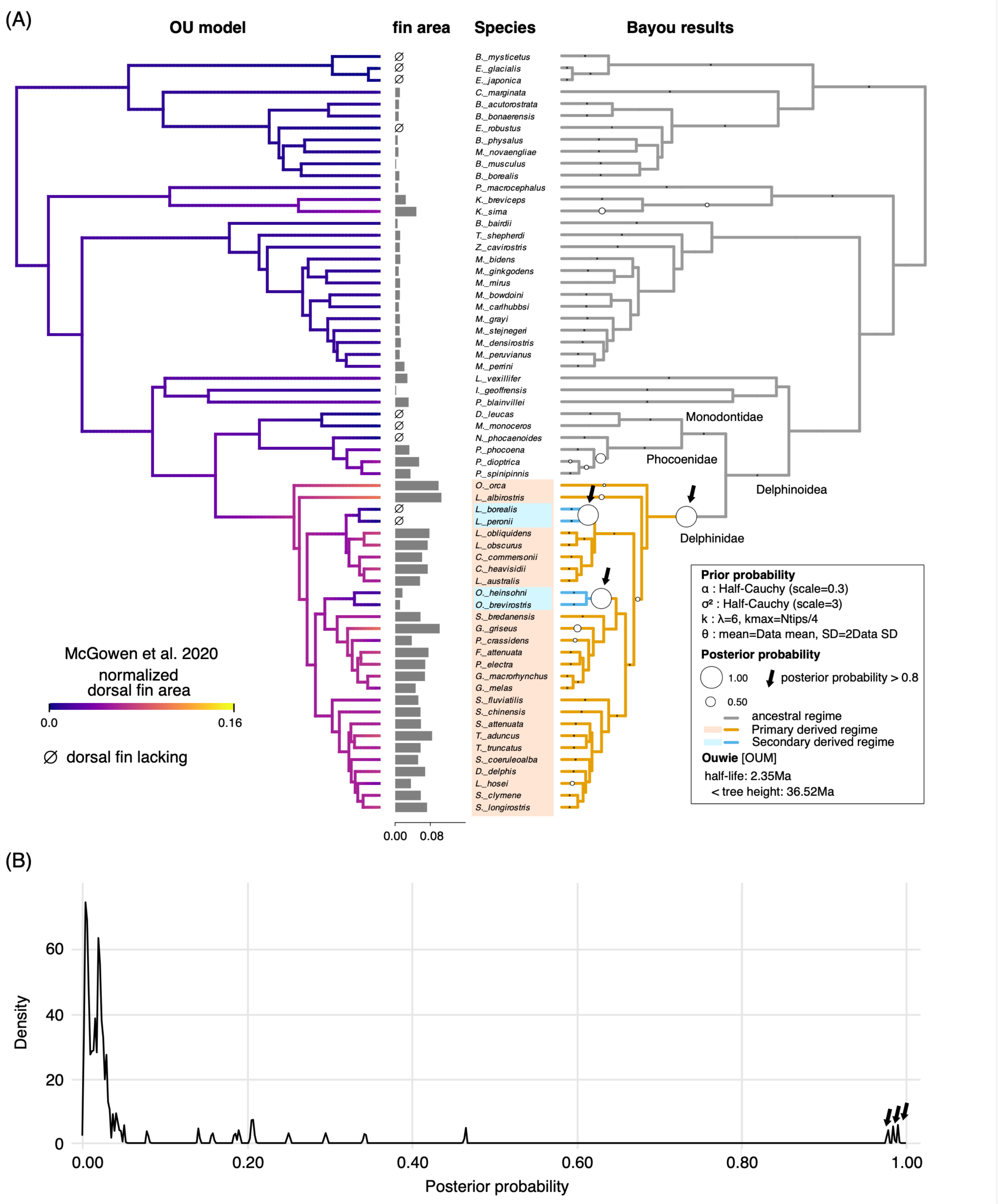


**Figure SC4 Ancestral state reconstruction and inferred evolutionary shifts in normalized dorsal fin area on the phylogeny of McGowen et al. (2020).**

(A) From left to right, the Ornstein–Uhlenbeck (OU) model ancestral state reconstruction, normalized dorsal fin area, species names, and Bayou posterior probabilities are shown. The legend also provides the numerical values of the priors used for each parameter. Arrows indicate branches with posterior probabilities >0.8, interpreted as significant evolutionary shifts. Species names are highlighted for groups inferred to have undergone such shifts. (B) Distribution of posterior probabilities across branches in Bayou. The horizontal axis shows posterior probability (0–1.0), and the vertical axis shows the number of branches. Arrows indicate branches with posterior probabilities >0.8, corresponding to the branches shown in (A).

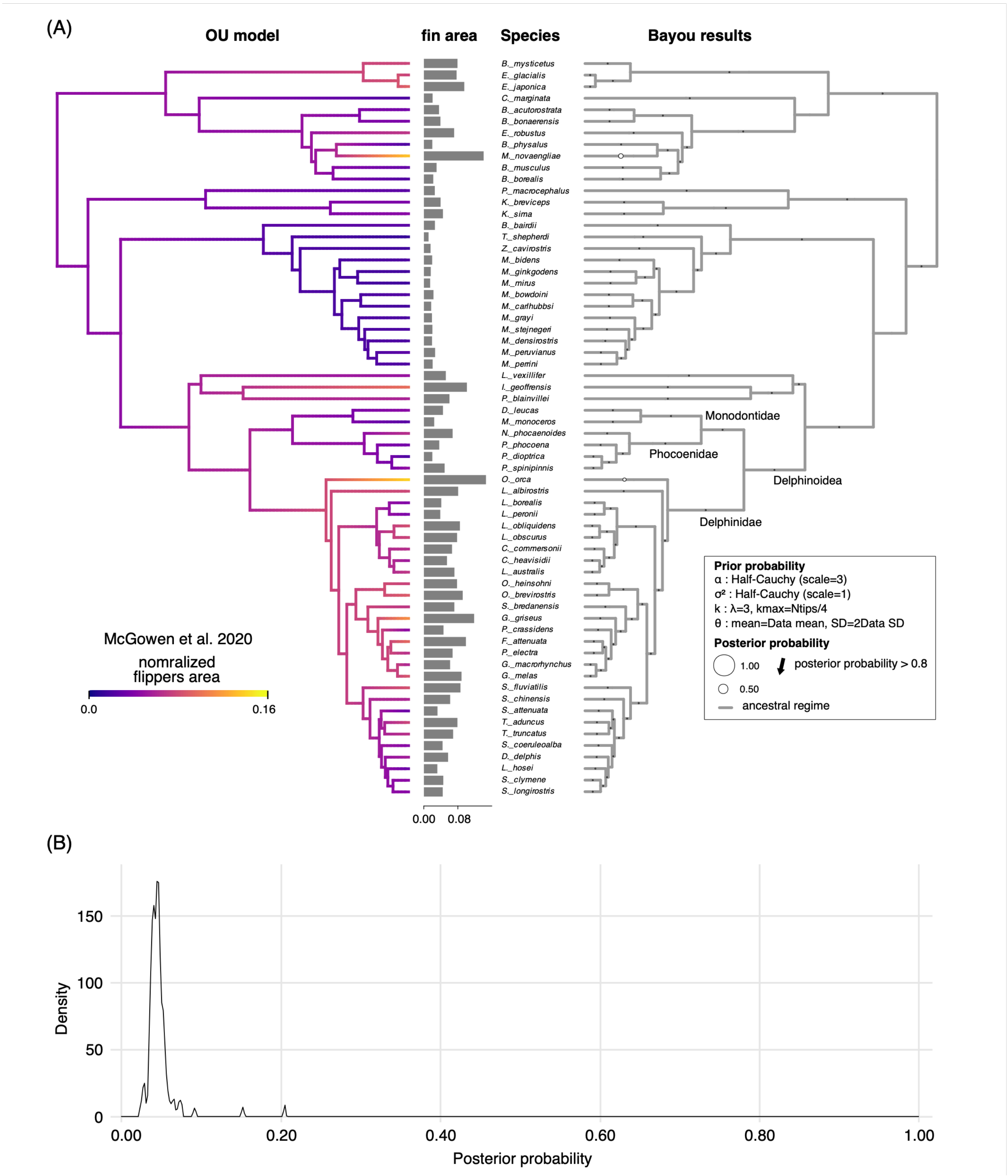


**Figure SC5 Ancestral state reconstruction and inferred evolutionary shifts in normalized flippers area on the phylogeny of McGowen et al. (2020).**

(A) From left to right, the OU model ancestral state reconstruction, normalized flippers area, species names, and Bayou posterior probabilities are shown. The legend also provides the numerical values of the priors used for each parameter. Arrows indicate branches with posterior probabilities >0.8, interpreted as significant evolutionary shifts. Species names are highlighted for groups inferred to have undergone such shifts. (B) Distribution of posterior probabilities across branches in Bayou. The horizontal axis shows posterior probability (0–1.0), and the vertical axis shows the number of branches. Arrows indicate branches with posterior probabilities >0.8, corresponding to the branches shown in (A).

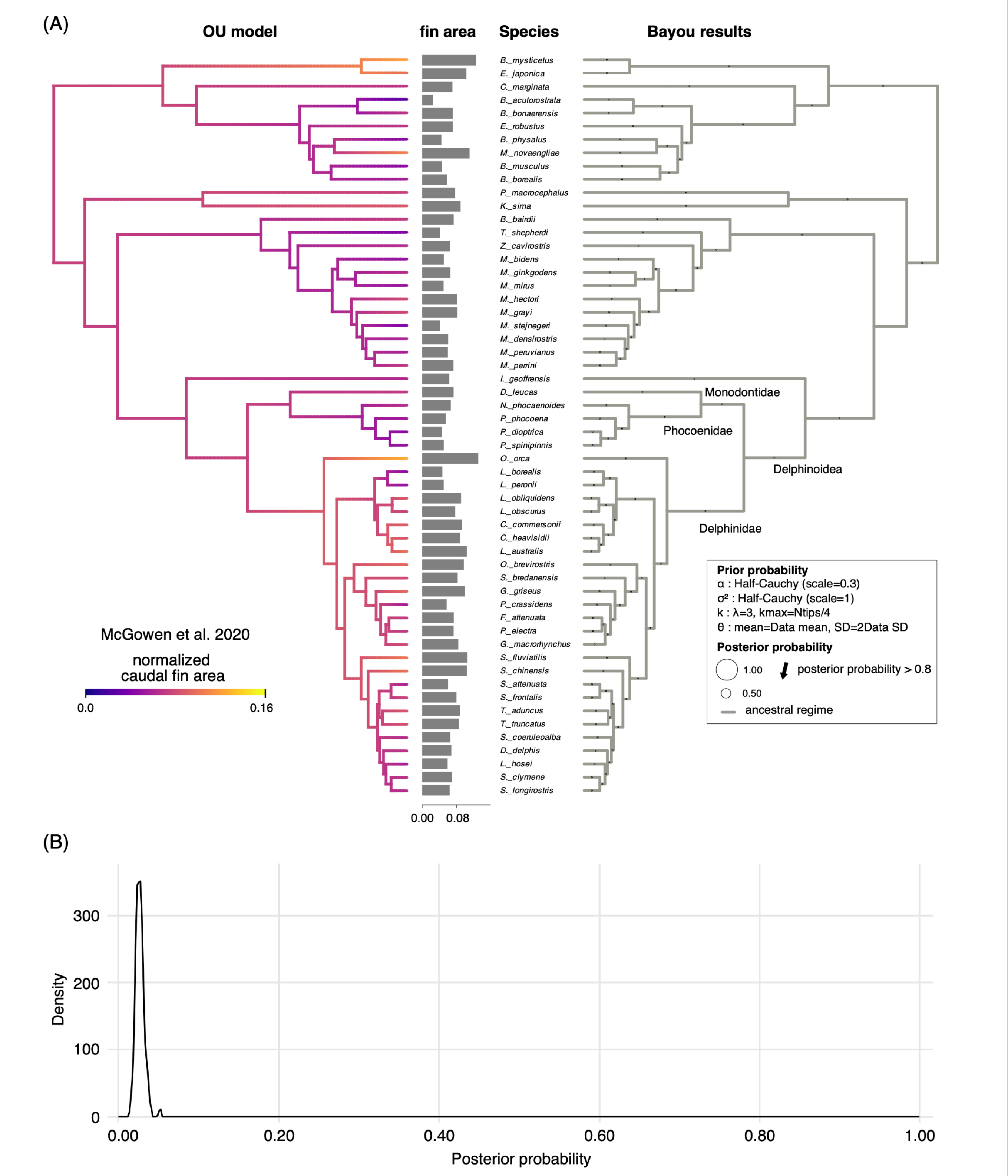


**Figure SC6 Ancestral state reconstruction and inferred evolutionary shifts in normalized caudal fin area on the phylogeny of McGowen et al. (2020).**
(A) From left to right, the OU model ancestral state reconstruction, normalized caudal fin area, species names, and Bayou posterior probabilities are shown. The legend also provides the numerical values of the priors used for each parameter. Arrows indicate branches with posterior probabilities >0.8, interpreted as significant evolutionary shifts. Species names are highlighted for groups inferred to have undergone such shifts. (B) Distribution of posterior probabilities across branches in Bayou. The horizontal axis shows posterior probability (0–1.0), and the vertical axis shows the number of branches. Arrows indicate branches with posterior probabilities >0.8, corresponding to the branches shown in (A).

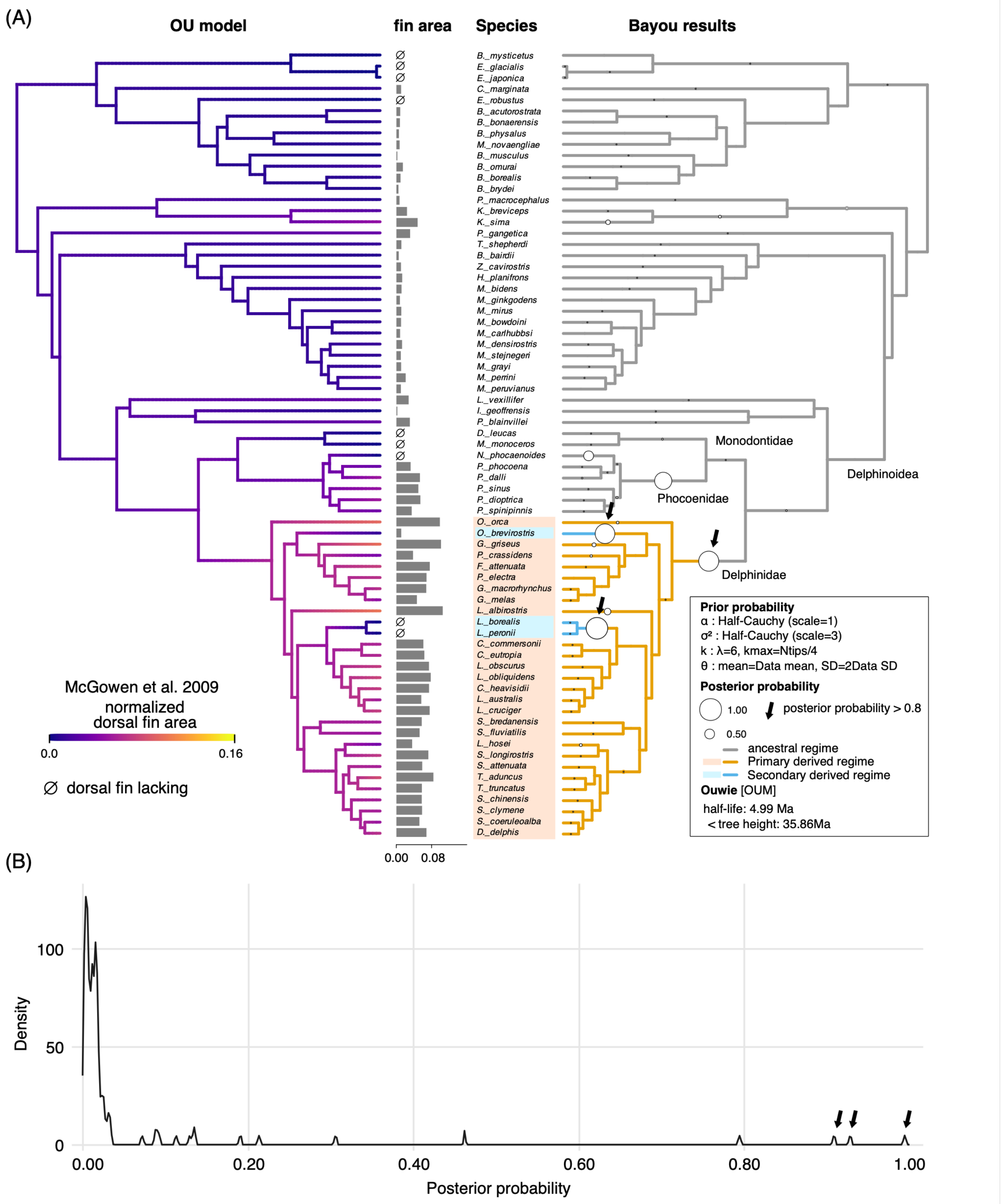


**Figure SC7 Ancestral state reconstruction and inferred evolutionary shifts in normalized dorsal fin area on the phylogeny of McGowen et al. (2009).**
(A) From left to right, the OU model ancestral state reconstruction, normalized dorsal fin area, species names, and Bayou posterior probabilities are shown. The legend also provides the numerical values of the priors used for each parameter. Arrows indicate branches with posterior probabilities >0.8, interpreted as significant evolutionary shifts. Species names are highlighted for groups inferred to have undergone such shifts. (B) Distribution of posterior probabilities across branches in Bayou. The horizontal axis shows posterior probability (0–1.0), and the vertical axis shows the number of branches. Arrows indicate branches with posterior probabilities >0.8, corresponding to the branches shown in (A).

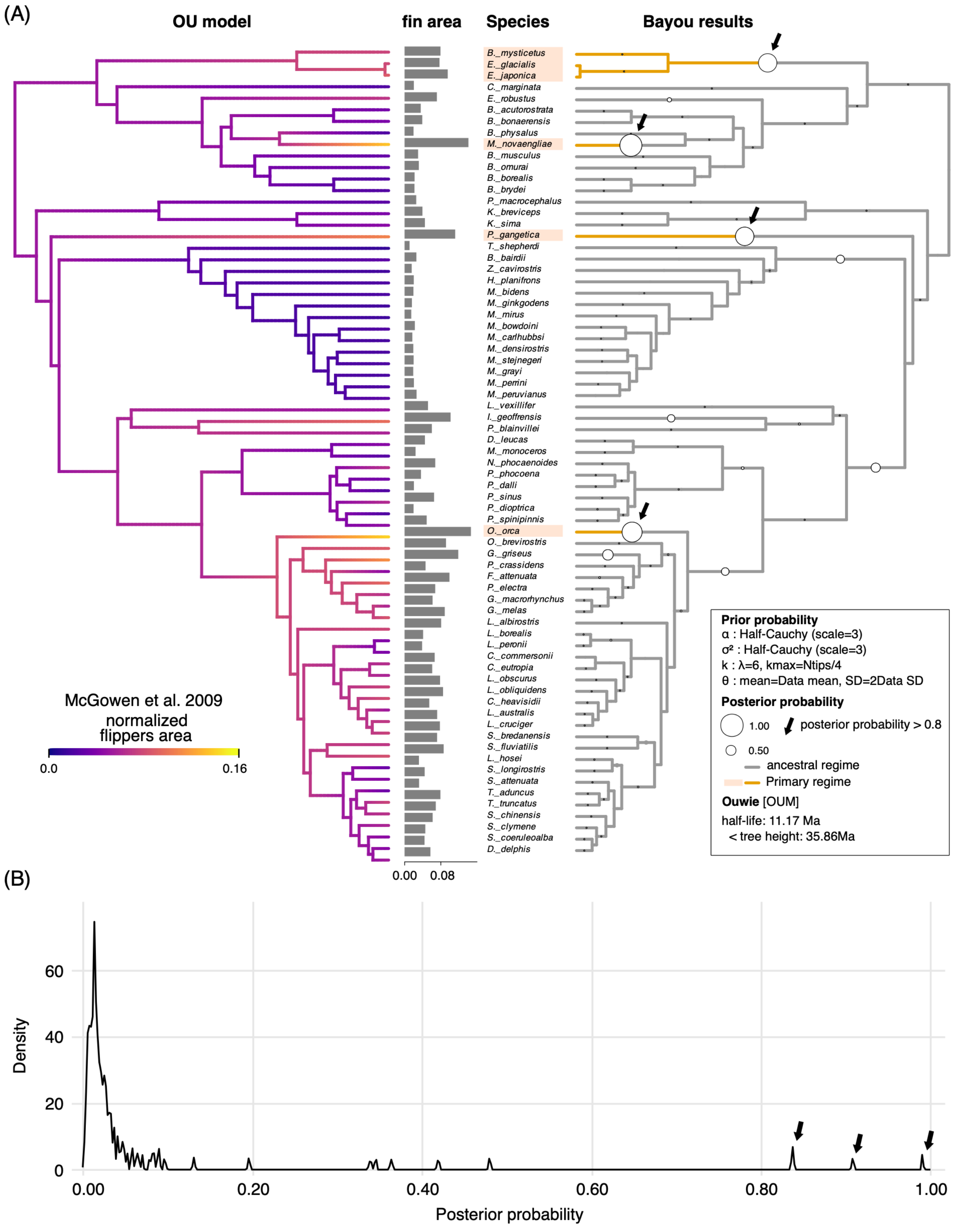


**Figure SC8 Ancestral state reconstruction and inferred evolutionary shifts in normalized flippers area on the phylogeny of McGowen et al. (2009).**

(A) From left to right, the OU model ancestral state reconstruction, normalized flippers area, species names, and Bayou posterior probabilities are shown. The legend also provides the numerical values of the priors used for each parameter. Arrows indicate branches with posterior probabilities >0.8, interpreted as significant evolutionary shifts. Species names are highlighted for groups inferred to have undergone such shifts. (B) Distribution of posterior probabilities across branches in Bayou. The horizontal axis shows posterior probability (0–1.0), and the vertical axis shows the number of branches. Arrows indicate branches with posterior probabilities >0.8, corresponding to the branches shown in (A).

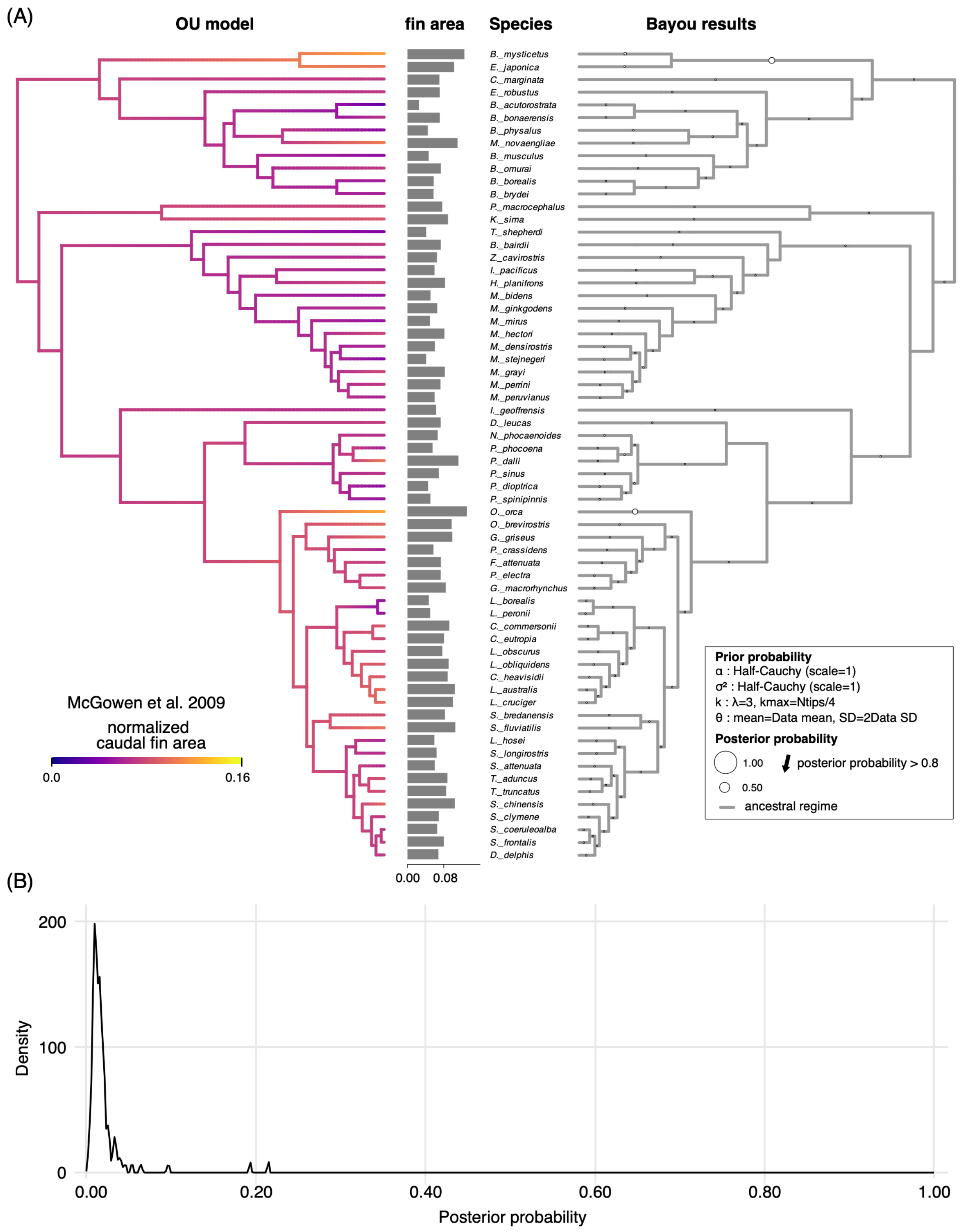


**Figure SC9 Ancestral state reconstruction and inferred evolutionary shifts in normalized caudal fin area on the phylogeny of McGowen et al. (2009).**

(A) From left to right, the OU model ancestral state reconstruction, normalized caudal fin area, species names, and Bayou posterior probabilities are shown. The legend also provides the numerical values of the priors used for each parameter. Arrows indicate branches with posterior probabilities >0.8, interpreted as significant evolutionary shifts. Species names are highlighted for groups inferred to have undergone such shifts. (B) Distribution of posterior probabilities across branches in Bayou. The horizontal axis shows posterior probability (0–1.0), and the vertical axis shows the number of branches. Arrows indicate branches with posterior probabilities >0.8, corresponding to the branches shown in (A).

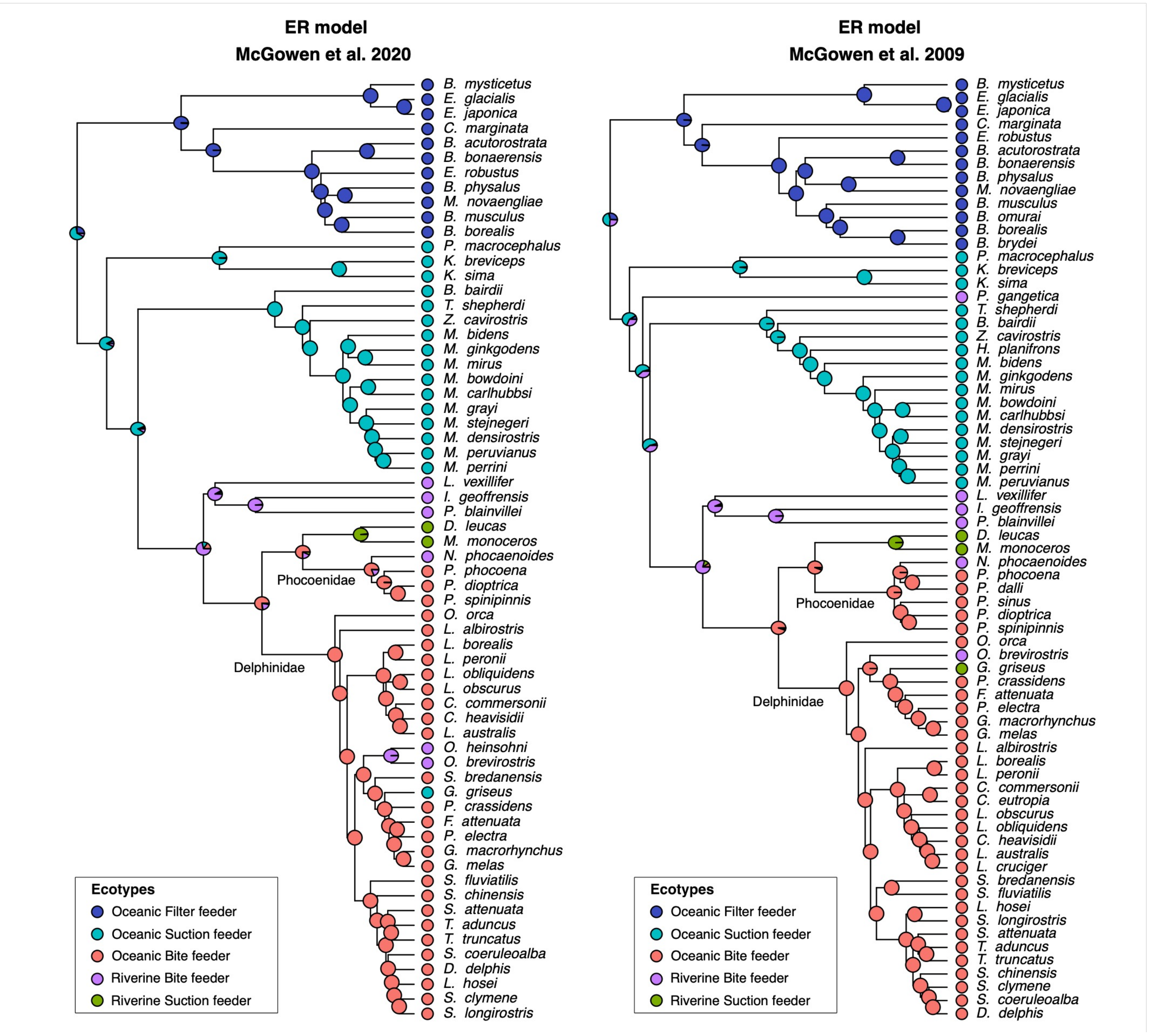


**Figure SC10 Ancestral state reconstruction of ecotypes.**

Ancestral state reconstruction of ecotypes under the ER model is shown for the molecular phylogenies of McGowen et al. (2020) and McGowen et al. (2009). Ecotypes were classified as oceanic filter feeder, oceanic suction feeder, oceanic bite feeder, riverine bite feeder, and riverine suction feeder. In both phylogenies, the common ancestor of extant cetaceans was not strongly reconstructed as any single ecotype, whereas each ecotype became strongly associated with particular lineages.
